# Tracking human foragers and their prey reveals adaptive predator–prey dynamics

**DOI:** 10.64898/2026.09.21.752541

**Authors:** Félicie Dhellemmes, Alexander Schakowski, Valerii Chirkov, Marwa M. Kavelaars, Fleur F. A. Korzilius, Dominik Deffner, Pawel Romanczuk, Raine Kortet, Pyry Pihlasvaara, Ralf H.J.M. Kurvers

## Abstract

Hunting for mobile prey is thought to have played a key role in hominin evolution, by providing high-quality nutrition that supported the development of the exceptionally large human brain (1; 2; 3; 4). However, human–prey dynamics remain poorly understood because studies have not yet tracked human foragers and their prey simultaneously. Here, we employ high resolution tracking of groups of human foragers (ice-fishers) and their prey (fish shoals) to study human–prey dynamics. Our results show that foragers adaptively combined personal and social information in deciding where to forage and for how long, closely matching the prey distribution. Prey responded dynamically to human exploitation, showing increased attraction to fishing activity, alongside decreased biting probability. Furthermore, we found that foragers adaptively relied on memory, preferentially returning to areas with high prey presence, particularly when their current return rate was low. Our results show how human foragers overcome the challenges of extracting invisible, mobile and reactive prey by tightly tuning patch-selection, patch-leaving and patch-return decisions to the distribution and behaviour of their prey.

## Main

Foraging for mobile prey is believed to have been a key driver of hominin evolution (1; 2). Humans and their hominin ancestors have consumed animal foods for over three million years (5; 6), with animal foods making up a relatively large proportion of their diet compared with their closest relatives (7). Access to animal proteins and nutrients is considered to have played a key role in the development of humans’ unusually large, energetically costly, and complex brains (3; 4). Moreover, targeting mobile prey presents exceptional cognitive and physical challenges, such as anticipation of both immediate and longer-term prey behaviour, use of complex tools, social coordination and high energy expenditure. The cognitive demands of overcoming these challenges may, in turn, have contributed to the evolution of human cognition (7; 8; 9). However, understanding of human–prey dynamics in situ remains limited because no previous studies have simultaneously tracked human foragers and their prey. From the predator perspective: How do human foragers integrate personal and social information to target mobile prey, and how adaptive are these strategies? How do they integrate information on previous prey encounters across time to strategically plan revisits? From the prey perspective: How do prey react to human predation, and how do alterations in prey behaviour feed back into human decision making?

Previous studies tracking human subsistence foragers (10; 11; 12) and participants in competitive foraging events (13; 14) have shown that humans strategically reduce relocation distance when they encounter potential food items, including animal prey (i.e., area-restricted search (15)). However, these studies did not simultaneously track the prey, leaving it unresolved how adaptive these search strategies were, and how the dynamics between humans and their prey unfold. Studies investigating the response of prey to human hunting have shown that hunting pressure typically reduces the spatial and exploratory behaviour of prey populations (16; 17). Moreover, human predation pressure over extended periods of time can lead to rapid changes in prey morphology and behaviour by selection (18; 19; 20). These studies show large-scale behavioural or evolutionary prey responses, but do not capture the fine-scale behavioural dynamics between humans and their prey.

In the non-human foraging literature, a long-standing research tradition has documented the decision-making strategies that enable foragers to optimize energy intake across heterogeneous resource landscapes (21; 22; 23). However, this research has largely focused on static, depletable resources; far less attention has been paid to the foraging context experienced by many predatory species, where the resource is itself mobile and reactive to the forager’s behaviour (24; 25).

To study the dynamics of human–prey interactions, we simultaneously tracked groups of human foragers and their mobile prey. We equipped groups of competitive ice-fishers with GPS tracking devices and head-mounted cameras, recording spatial search and foraging success. Simultaneously, we used multiple real-time sonars to track their prey, mobile fish shoals, under the ice. Combining high-resolution tracking of humans and their prey with computational modeling techniques, we show how humans integrate personal information (i.e., their own catching success) with social information (i.e., the presence of others) across space and time to target mobile prey, and test the adaptiveness of these strategies. Tracking prey at both angling spots and unexploited areas, we quantify the prey’s spatial (e.g., redistribution) and behavioural (e.g., biting likelihood) responses to exploitation. Finally, we show how humans react to dynamic prey behaviour over time by preferentially remembering and returning to successful foraging spots.

### Tracking human foragers and their prey

We organized 12 ice-fishing competitions at three lakes in Northern Karelia, Finland (15–22 March 2024; Extended Data Fig. 1). Ice-fishing is a traditional practice in the Nordic countries and was once an important means of subsistence (26). In modern times, regional and national ice-fishing competitions are organized across these regions, keeping the practice alive (27). At each lake, we created four arenas of approx. 7,000m^2^, with every competition taking place in a different arena. In each competition, six highly experienced ice-fishers (mean years of experience = 53, standard deviation (SD) = 8) competed against each other for two hours. They were free to drill, fish and relocate within the arena. At the end of each competition, each participant’s total catch (of Eurasian perch, *Perca fluviatis*) was weighted and participants were reimbursed based on their total catch weight (see Methods for incentivization). Each participant was equipped with a GPS device logging high-resolution spatial coordinates data and with a head camera filming continuously (Fig. 1A). The head camera recordings were manually annotated; for each spot visited, we recorded when participants started drilling, started fishing, caught fish and stopped fishing by leaving the spot. During a competition, participants walked, on average, 774m (SD across individuals = 285; Extended Data Fig. 2A) to visit 26.3 (SD = 9.3) angling spots, most of which were unsuccessful (i.e., no fish were caught, mean = 20.2 *±* 8.9 SD, Extended Data Fig. 2B). This resulted in a total of 1,891 angling spot choices across all competitions. Participants spent, on average, 31.7s (SD = 23.9; Extended Data Fig. 2C) relocating between subsequent angling spots, 27.2s (SD = 9.3; Extended Data Fig. 2D) drilling per angling spot, 124.4s (SD = 108.6; Extended Data Fig. 2E) fishing at unsuccessful angling spots, and 530.2s (SD = 540.9; Extended Data Fig. 2F) fishing at successful angling spots (i.e., spots at which they caught at least one fish). Participants caught, on average, 28.3 fish per competition (SD = 21.7; Extended Data Fig. 2G). See Supplementary Table 1 for details per competition.

**Fig. 1.**
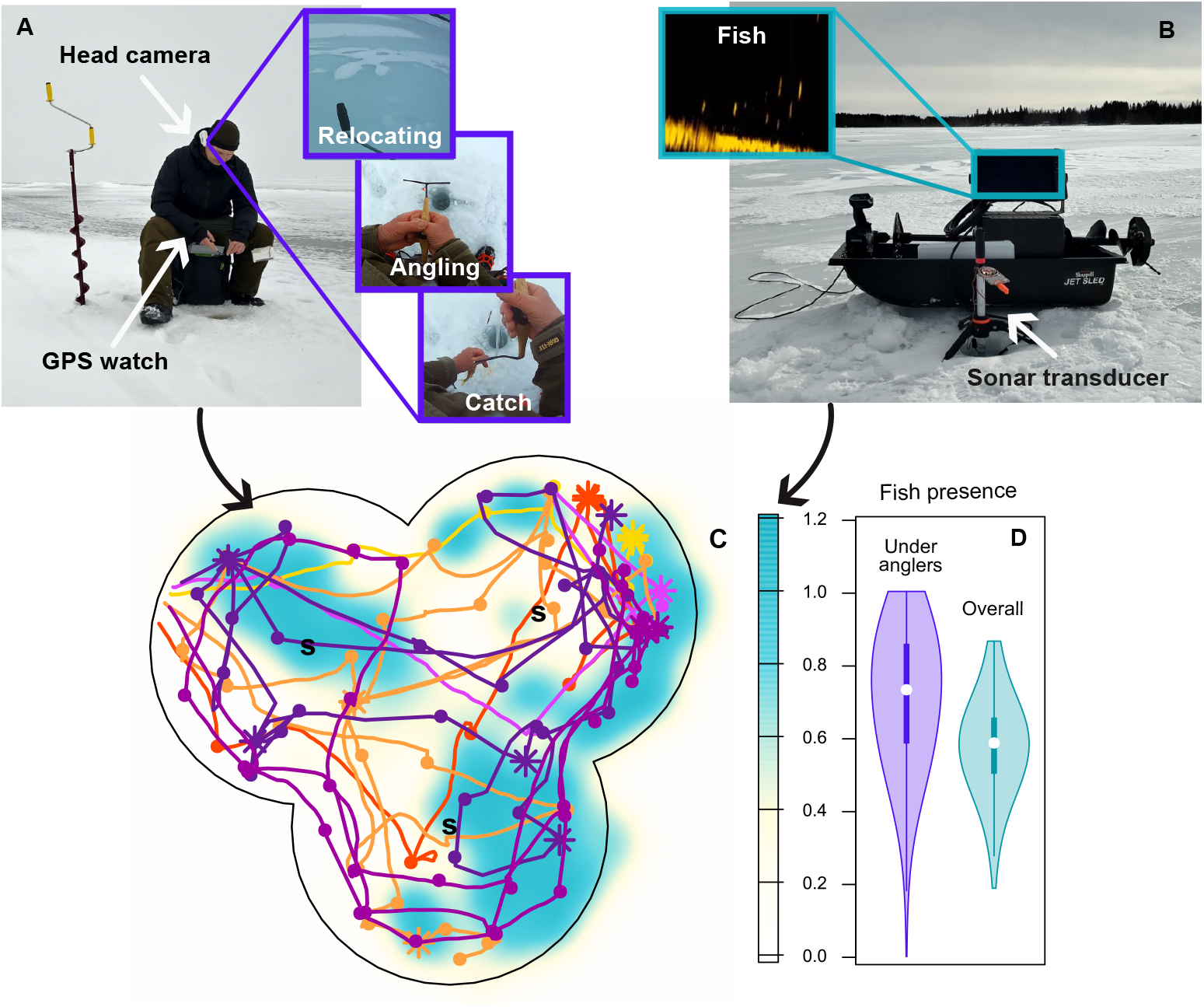
Human and fish tracking methods. (A) Participant fishing (Pictured is author **PP** competing in a different event; Image copyright: **FD**). Participants were equipped with GPS smart-watches and head cameras that provided continuous data on two behavioural states (relocating and angling) and on fish catches. (B) Sonar set-up (Image copyright: **FD**). The transducer was placed below the ice using a tripod; the fish were visible on an external monitor. (C) GPS tracks of participants combined with the fish presence estimated from the sonar data. The spatial trajectories of six humans are shown as lines connecting successful (i.e., at least one fish was caught; stars) and unsuccessful (i.e., no fish was caught; dots) spot visits. Fish presence is shown in a white–to–blue gradient for one minute of the competition. The arena boundary is shown in black and was created to ensure that participants remained within 30m of a sonar. s = sonar locations. (D) Fish presence at active angling spots was higher than the average fish presence in the arena (Estimate [95% credible interval] = *−*0.13[*−*0.14*, −*0.13], see Extended Data Fig. 3A, B).

We simultaneously monitored the fish presence under the ice using three sonars (Lowrance ActiveTarget2 Live Sonar) placed equidistant of each other in a triangular position, ensuring complete coverage of the arena (Fig. 1B; Extended Data Fig. 1B–D). During the competitions, the sonars’ transducers were lowered under the ice and continuously rotated (angle of the sonar beam: 18°). When fish appeared on the sonar display, we recorded the time and the bearing and distance of the fish relative to the transducer, and we estimated the number of fish in the shoal and its spatial dimension (see Methods for details). Once the transducer made a full 360° circle, we rotated it back and started a new rotation. The average time per rotation was 3.7min (SD = 3.3; Extended Data Fig. 2H) and the average number of detected fish per rotation was 18.3 (SD = 30.9; Extended Data Fig. 2I).

For each encountered fish shoal, we calculated the GPS coordinates of the center and edges of the shoal and estimated the number of fish in the shoal. We then randomly generated the corresponding number of point coordinates within the shoal’s outer edges, providing a single coordinate for each fish. Because the sonar beam passes every location every 3.7 minutes on average, we interpolated these data in three dimensions (northing, easting and time) using Gaussian Process (GP) models to create a continuous intensity surface of fish presence through space and time (Fig. 1C). We first ran hyperparameter optimization for each competition and then fitted GP models on the fish coordinates for each competition using the optimized hyperparameters. The fitted models were used to predict fish presence every minute in 1m² grid cells across the entire arena. This approach generated a dynamic spatiotemporal surface of predicted fish presence in which areas close to (far away from) a fish detection in space and time retained high (low) fish presence. Across competitions, the optimized hyperparameters yielded an average length scale parameter of 6m (SD = 0.9) for the spatial kernel and 16.4 minutes (SD = 0.6) for the temporal kernel. On average 40% of grid cells had a fish presence above 0.8 (SD = 10%, Extended Data Fig. 2J).

Combining the spatial data of humans and fish with catch information (Fig. 1C), we found that the estimated fish presence at successful spots (i.e., locations where individuals caught at least one fish) was higher than at unsuccessful spots (i.e., locations where individuals did not catch fish) (Extended Data Fig. 3C). Importantly, participants selectively fished at locations with above-average fish presence relative to the rest of the arena (Fig. 1D, Extended Data Fig. 3A; see Extended Data Fig. 3D–F for results per lake), resulting in an overlap between areas of high fish presence and participant density. In the following, we unpack the decision-making mechanisms leading to the adaptive exploitation of this mobile, invisible prey.

### Adaptive personal and social information use for spot choice

We first examined how participants decided where to go. Overall, participants chose spots with a higher fish presence on arrival than expected by chance (Extended Data Fig. 3G). To investigate how they achieved this, we analyzed which information features participants used to select their next spot, and how well these features mapped onto the actual fish distribution under the ice.

To disentangle the features driving participants’ spot choices, we extended a modeling framework introduced by (13), which consists of a step-selection model commonly used to understand animal movement decisions (28). For each spot choice made by a participant, we generated 30 alternative spots in the vicinity and compared the features of the chosen spot against the alternative spots. For each spot, we calculated the following set of features that may drive spot selection: (i) social feature: the spatial density of other participants; (ii) success feature: the spatial density of previous successful spots visited by the participant; (iii) loss feature: the spatial density of previous unsuccessful spots visited by the participant. We also included the distance to the closest sonar in the model to account for sonar presence in the arena.

For the social, success and loss features, we also estimated a spatial bandwidth parameter, which governs the rate of spatial decay. Low bandwidth estimates indicate that individuals rely on a feature only in its immediate vicinity, while high bandwidth estimates indicate that they generalize a feature across larger distances.

For the success and loss features, we also estimated a temporal bandwidth parameter, which describes how the influence of a feature decreases across time. To avoid confounding with the time participants spent at a spot, we modeled this decay as a function of the number of spots visited since. High temporal bandwidth estimates indicate that the influence of a previously visited successful (unsuccessful) spot persists across many subsequent relocations, whereas low temporal bandwidth estimates indicate that the effect of previously visited locations on movement is ephemeral.

Spatial and temporal feature weights combined with bandwidth parameters allowed us to extrapolate individuals’ mental representations of the resource distribution across time and space (29; 30). The model included offsets per trip (a person within a competition), per participant and per competition (see Methods for full details).

Participants used both their own catching success (i.e., personal information) and the location of others (i.e., social information) to select their next angling spot (Fig. 2A, Supplementary Table 2). Participants were more likely to select spots that were close to other participants, close to previously successful spots, and further away from previously unsuccessful spots. Participants also preferred fishing far away from the sonars (Extended Data Fig. 4, Supplementary Table 2), possibly to reduce competition by targeting fish outside of the arena’s boundaries.

**Fig. 2.**
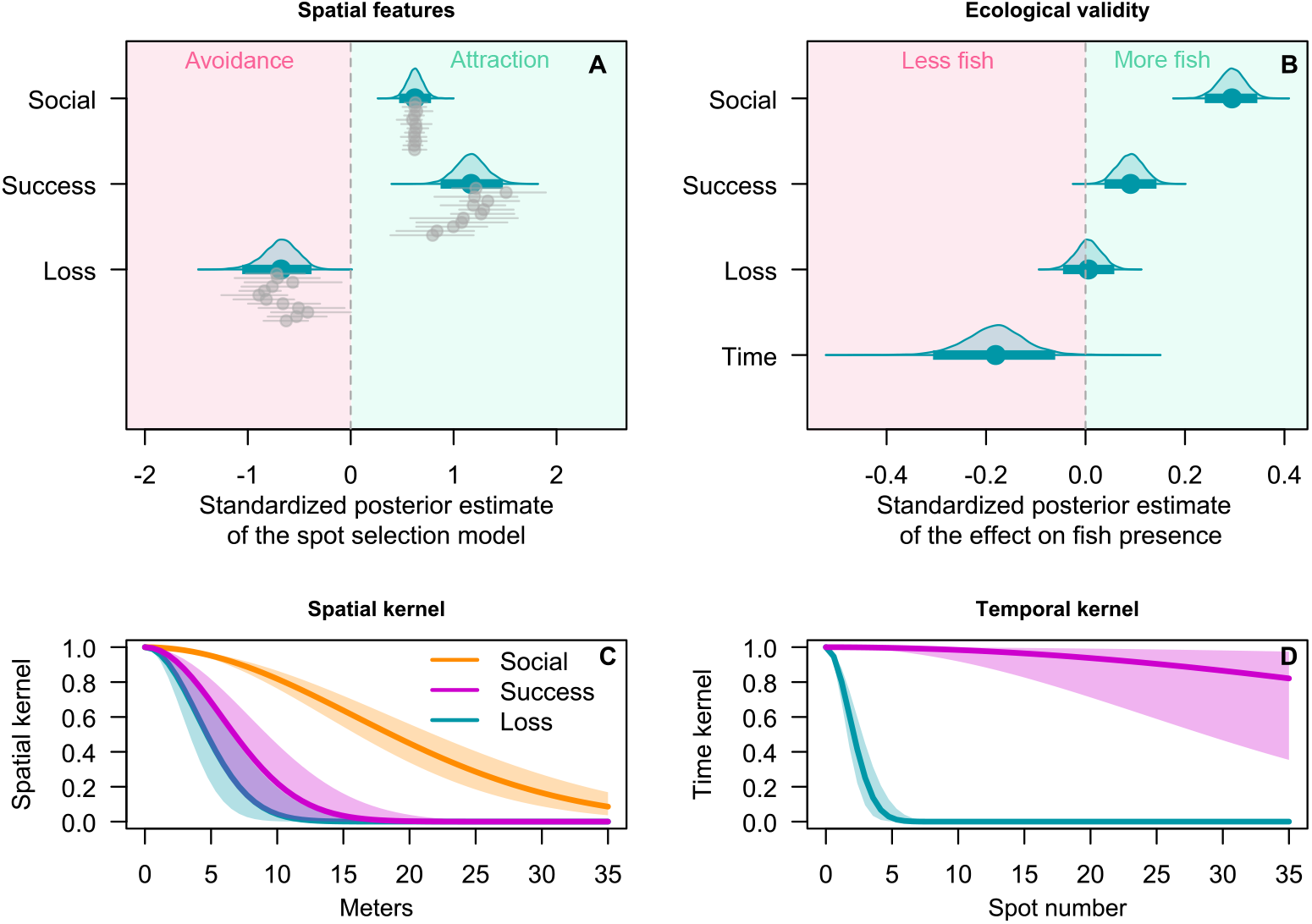
Results of the spot selection model. (A) Standardized posterior estimates of the spot selection model. Positive values indicate attraction to a feature; negative values indicate avoidance. Estimates are standardized such that a one unit increase corresponds to a one SD increase in the respective feature. Blue dots and lines show the overall point estimates and 95% posterior credible intervals. Grey dots and lines show the per-competition point estimates and 95% posterior credible intervals. (B) Ecological validation of the spot selection model showing how predictive each feature is of fish presence. All variables are z-standardized. (C) Posterior predictions of spatial feature generalization for personal and social information. (D) Posterior predictions of temporal feature generalization for personal information. Kernels are calculated using the model’s bandwidth estimates. Ribbons indicate the 95% credibility intervals. See Supplementary Table 2 for the posterior estimates of the spot selection model and Supplementary Table 3 for the posterior estimates of the ecological validation analysis.

To determine how these features mapped onto fish presence—in other words, whether the anglers’ strategies were aligned with where fish actually were—we predicted the fish presence measured at a chosen spot (computed as the average fish presence in a 3m radius at the time of spot choice) from the value of the social, success and loss feature at the time of spot choice and the distance to the sonar. We also included time in the competition to control for possible fish depletion or attraction over time. Participants’ use of the spatial features was largely in line with fish presence: Chosen spots close to other participants and close to previously successful spots had higher fish presence than spots further away from other participants and from previously successful spots (Fig. 2B, Supplementary Table 3). In contrast, there was no difference in fish presence between spots close to or further away from previously unsuccessful spots. Fish presence decreased with distance to the sonar ( Supplementary Table 3), most likely due to measurement error, as the likelihood of an object being detected decreases with distance to a sonar (31). Additionally, fish presence decreased over the course of the competition (Extended Data Fig. 3B, Supplementary Table 3), suggesting local depletion.

Participants integrated personal and social information at different spatial scales (Fig. 2C): Social information was generalized over a larger spatial distance than personal successes and losses, suggesting that it is treated as spatially less precise.

Interestingly, participants also integrated successes and losses at different temporal scales (Fig. 2D). The effect of a previous loss (i.e., an unsuccessful spot visit) decayed quickly over subsequent spot choices. In contrast, a success (i.e., a successful spot visit) decayed slowly: Participants remained attracted to their previously successful spots throughout the competition, suggesting that they continued to consider them areas of interest rather than completely fished out. We return to the temporal development of fish presence at (un)successful spots and the dynamics driving spot revisits later. First, we examine how participants decided when to leave a spot.

### Adaptive personal and social information use for spot leaving

We next studied how participants decided when to leave a spot and how adaptive these decisions were. Overall, participants stayed longer at spots with higher fish presence upon arrival (Extended Data Fig. 5A). To determine how they achieved this, we studied which spatial and temporal features governed decisions to leave a spot. For every 10-second interval spent fishing, we classified whether the participant left or not. We tested the same features as in the spot selection model: proximity to previous successful and unsuccessful spots, density of competitors (using the previously estimated bandwidth parameters) and sonar distance. We also included whether or not a participant caught a fish at the spot and the time since the last catch there (or since arrival at the spot in case of no catch). We also included fish presence in the model as participants may get additional sensory information via the fishing line (e.g., nibbles without a catch). The model offsets were the same as for spot leaving (see Methods for full details).

The strongest predictor of participants’ likelihood of leaving was whether or not they caught a fish, with participants being less likely to leave after catching a first fish (Fig. 3A, Supplementary Table 4). The longer participants did not catch a fish, the more likely they were to leave. Participants were also less likely to leave spots close to competitors. Proximity to previously successful or unsuccessful spots had no credible effect on the leaving likelihood. Interestingly, fish presence credibly reduced participants’ likelihood of leaving, suggesting that our features did not capture all information available to participants. Participants probably get more sensory information on fish presence via subtle movements of their fishing line. When relevant predictors (e.g., fish catch and social density) were removed from the model, the effect of fish presence increased (Extended Data Fig. 5B), further corroborating that the spatial and temporal features act as informational cues for participants to infer fish presence under the ice. Finally, participants were less likely to leave spots further away from the sonar (Supplementary Table 4), again suggesting that they were attracted to the arena boundaries.

**Fig. 3.**
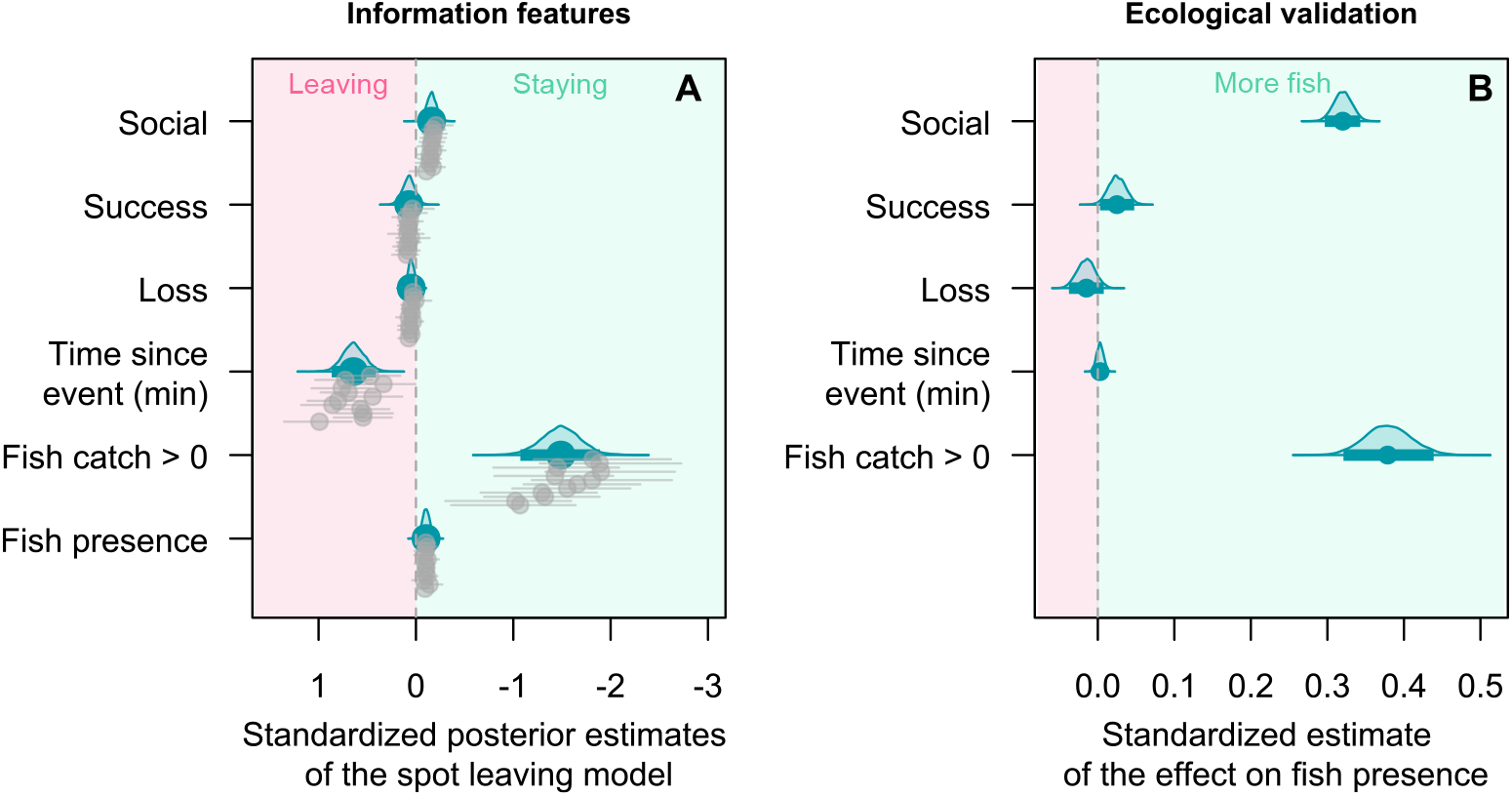
Results of the spot leaving model. (A) Standardized posterior estimates of the spot leaving model. The x-axis is inverted to facilitate interpretation. Estimates apart from time and catch are standardized such that a one unit increase corresponds to a one SD increase in the respective feature. Blue dots and thick lines show the overall point estimates and the 95% credible intervals. Grey dots and lines show point estimates and 95% credible intervals per competition. (B) Ecological validation of the spot leaving model showing how predictive each model feature is of fish presence. Social, success and loss features are z-standardized. See Supplementary Table 4 for posterior estimates of the spot leaving analysis and Supplementary Table 5 for posterior estimates of the ecological validation analysis.

To determine whether participants’ feature use was in line with fish presence, we developed a similar ecological validation model as for spot selection. For every minute, we computed the value of all spatial and temporal features (excluding fish presence) and modeled how well they predicted fish presence (estimated per minute). Three of the five features predicted fish presence in a manner consistent with how participants used them (Fig. 3B): Fish presence was higher at spots where fish were indeed caught and at spots with high social density, whereas proximity to previous losses did not predict fish presence (Fig. 3B, Supplementary Table 5). Proximity to previously successful spots had a small positive effect on fish presence (mean [95% credible intervals]: 0.03[0.00 *−* 0.05]), although this feature did not inform participants’ leaving decision. The most notable discrepancy was that the time since catch did not predict fish presence, despite participants consistently relying on it. Finally, spots further away from the sonar had lower fish presence, due to reduced detection likelihood by the sonar (Supplementary Table 5). Overall, participants’ use of temporal and spatial leaving cues corresponded well with the ecological validity of these cues in predicting fish presence, explaining their tendency to stay longer at spots with higher fish presence upon arrival (Extended Data Fig. 5A, Supplementary Table 5).

### Successful angling increases fish presence but reduces catches

We now turn to prey dynamics, showing how the fish respond spatially and behaviourally to angling. Being a mobile resource, fish can react spatially by moving away from areas with fishing pressure (32), but they can also be attracted to it (33). To test these possibilities, we compared the temporal development of fish presence at (un)successful spots to non-fished control spots. This approach allowed us to test how fish presence changes locally in response to active angling while accounting for global changes in fish dynamics in the arena. For each (un)successful spot visit, we identified up to five control spots (mean number of control spots = 3.6, median = 4, min = 1, max = 5) that were close in time to the (un)successful spot (within the same minute), had similar fish presence at the start of the angling event (difference smaller than 0.02), and had no participants within 10m (see Methods for details). We then determined the difference in fish presence over time relative to the start of the visit (t = 0) for both visited and control spots. Positive (negative) values indicate that fish presence increases (decreases) over time. We then modeled how fish presence developed at (un)successful spots relative to control spots.

At unsuccessful spots, fish presence decreased relative to control spots (Fig. 4A, Supplementary Table 6), suggesting that drilling and fishing cause local disturbances that deter fish (32). Conversely, successful spots showed increasing fish presence relative to control spots (Fig. 4A, Supplementary Table 7), possibly because fish actively inspect hooked conspecifics or are attracted to the smell of repeated baiting (participants rebaited after each capture (33)).

**Fig. 4.**
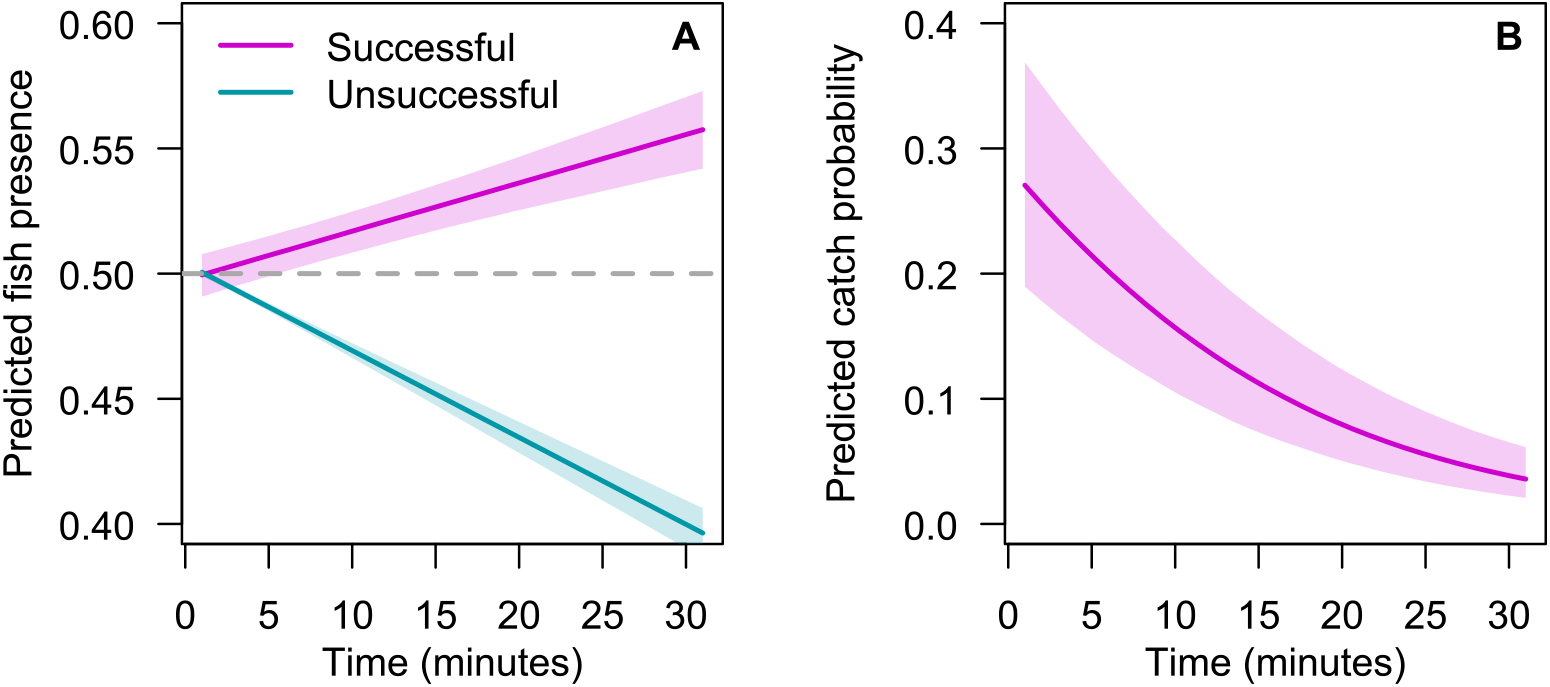
Impact of angling on fish presence and fish catches. (A) Predicted change in fish presence over time at successful and unsuccessful spots, relative to non-fished control spots. These results remained stable across different fish presence thresholds for control spot selection (Extended Data Fig. 6A, B). (B) Predicted catch probability over time at successful spots. See Supplementary Tables 6, 7 and 8 for posterior estimates.

Fish may also adjust their vigilance and willingness to bite under predation risk (17; 34). To test how the catch probability changed over the course of a spot visit, we modeled how fish presence and time since start of a spot visit affected catch probability at successful spots. Catch probability decreased over the course of a spot visit (Fig. 4B, Supplementary Table 8). This is striking because fish presence at successful spots increased gradually over the same period (relative to control spots, Fig. 4A). The combination of increasing fish presence and decreasing catch probability suggests a reduced likelihood of individual fish biting over time, possibly due to social contagion of predation risk (35). Supporting this explanation, longer intervals between a spot’s abandonment and its revisit were associated with higher catch probability upon revisit (Extended Data Fig. 6C, Extended Data Fig. 7, Supplementary Table 9, and see Methods for details), suggesting that fish return toward baseline biting levels when left undisturbed.

### Integrating past and current return rates for revisits

So far, the results show that human foragers exert a direct influence on the spatial distribution and behaviour of their prey. These prey dynamics, in turn, have important consequences for the strategies that participants should use to maximize resource returns. At successful spots, participants experienced decreasing capture rates over time (Fig. 4B). If this was due to local depletion, returning to these spots may not be adaptive. However, if fish densities remain high and baseline biting levels recover over time—as was observed in this study system—it may be adaptive to memorize these spots and revisit them. The highly experienced ice-fishers in our study may have learned to exploit these prey dynamics and strategically revisit previously successful spots. Our spot selection analysis already showed that successful spots indeed remain attractive throughout the entire competition (Fig. 2D). Here, we investigate the dynamics of spot revisits.

We identified spatial clusters of spots by applying a density-based spatial clustering of applications with noise (DBSCAN) algorithm to the sequence of spots visited by a participant. We used a maximum distance value (*ɛ*) of 10m (Fig. 5A), with spots within 10m of a previously visited spot being assigned to the same cluster and spots more than 10m away being assigned to a new cluster. Revisits were defined as a participant leaving a cluster and returning to it later; continuous exploration of the same area did not qualify as a revisit (Fig. 5A). When a cluster was revisited, all spots within it were labeled as revisited (see Methods for details). Participants made, on average, three cluster revisits during a competition (mean = 3.2 *±* 2.6 SD; Extended Data Fig. 8). In line with the results of our spot selection analysis, they were more likely to revisit previously successful than unsuccessful spots (Fig. 5B, Supplementary Table 10). Likewise, they were more likely to revisit spots with higher fish presence (measured at the point of departure from that spot, Fig. 5C, Supplementary Table 11), showing that revisits were selectively made to areas with more fish. Spots discovered later in the competition were less likely to be revisited (Supplementary Tables 10, 11).

**Fig. 5.**
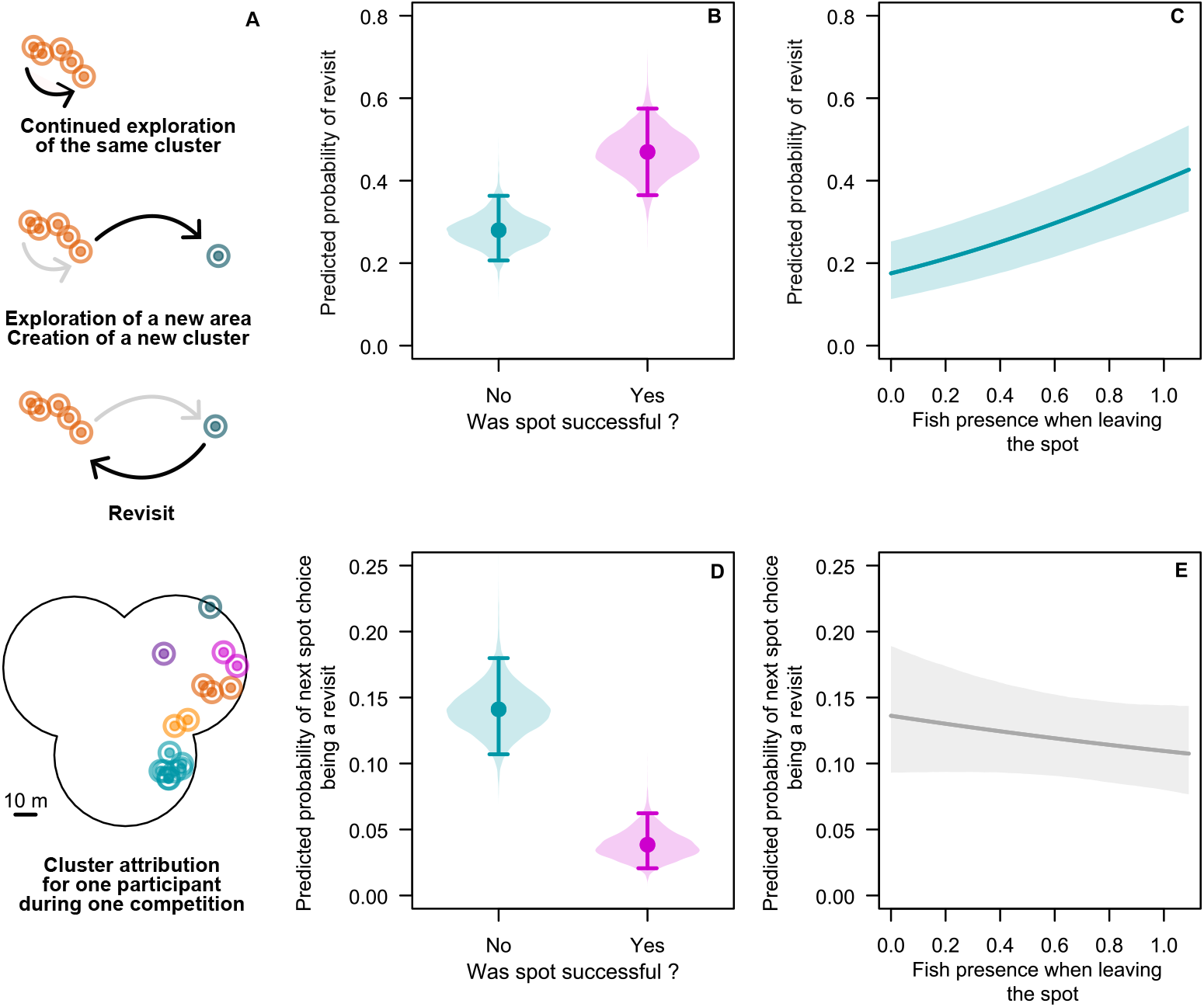
Results of the revisit analysis. (A) When relocating, participants could either remain in the same cluster, explore a new area creating a new cluster, or revisit a previous cluster. Each circle represents one spot, with same-coloured spots belonging to one cluster (with the outer circles representing the 10m threshold). Bottom image shows the cluster attribution of all spots for one foraging trip. (B,C) The probability that a spot (i.e., the cluster to which it belongs) is revisited in the future as a function of (B) whether or not a participant was successful in that spot and (C) fish presence when the participant left the spot. (D,E) The probability that the next spot is a revisit to a previous cluster as a function of (D) whether or not the last spot was successful and (E) fish presence at the last spot when the participant left that spot. See Supplementary Tables 10, 11, 12, 13 for the posteriors of the revisit model parameters.

Foraging theory predicts that the likelihood of revisiting a patch will depend not only on the resource value of that patch, but also on how it compares with other patches in the environment (36; 37). Indeed, participants were more likely to revisit a cluster directly after visiting an unsuccessful spot than a successful spot (Fig. 5D, Supplementary Table 12). Fish presence at the last spot showed a negative trend in predicting whether the next spot was a revisit to a previous cluster, though this effect was not credible (Fig. 5E, Supplementary Table 13). All results were robust across different values of the clustering parameter *ɛ* (Extended Data Fig. 6D–G). In sum, participants were more likely to revisit a cluster in which they were successful before and, by extension, clusters with a relatively high fish presence. Additionally, they were more likely to revisit a cluster after visiting an unsuccessful spot.

## Discussion

To our knowledge, this is the first study to simultaneously track human foragers and their prey in the wild. Our work shows that ice-fishers adaptively integrate personal and social information sources to locate areas with high fish densities and that they fish for longer in these areas. In addition, it shows that fish respond to successful angling by being attracted to it, but also reducing their willingness to bite. Finally, our findings show how humans use prey recovery by strategically revisiting previously successful areas.

Our expert foragers used three distinct mechanisms to selectively choose spots with high fish density. First, they used area-restricted search—a well-known search mechanism across species and contexts (15; 38)—by temporarily avoiding areas close to unsuccessful spots, while seeking out areas close to successful spots. In other words, they made relatively short relocations when encountering fish, and relatively long ones otherwise, in line with findings of previous human foraging studies (11; 12; 13; 14; 39). Second, they strategically revisited spots with high fish presence later in the competition. Whereas previously unsuccessful spots affected spot choice only briefly, previously successful spots remained attractive throughout the competition. This finding suggests that humans may not store the location of previously unsuccessful locations in memory for long, but rather use their limited memory capacity (40) to store previously successful locations. Given that our foragers predominantly encountered unsuccessful spots (Extended Data Fig. 2B), remembering the few highly valuable locations constitutes an efficient strategy, in line with adaptive memory processing (41). It is possible that fish catches increase arousal, which can help consolidate memory storage (42). Third, foragers were attracted to areas where others were fishing, and social density positively predicted fish presence. The role of the social environment remains understudied in human behavioural ecology, with most previous field studies focusing on solitary foragers (11; 12; 37). Our findings address recent calls to dedicate more attention to the role of social components in human foraging (43; 44), demonstrating that social information is integral to foraging success—a pattern consistent with the broader animal kingdom (45),

Participants also fished for longer at spots with high fish presence through four mechanisms. First, they were less likely to leave a spot after a first catch, in line with previous findings in human search (13; 46). Second, they remained longer at spots close to competitors. Third, fish presence directly reduced participants’ leaving likelihood, suggesting that they were able to use information not captured by our features to infer fish presence (e.g., tactile information from the fishing line). Fourth, the longer they did not catch anything, the more likely participants were more to leave a spot. Such giving-up time rules are predicted to outperform more static rules across a range of resource environments (46; 47). However, time since last catch did not predict fish presence at a spot, possibly because fish presence is an imperfect measure of what matters directly to a forager, namely, the average return rate. Because prey respond to exploitation in complex ways, fish presence, though an important predictor of catch success, is an incomplete one.

Our work reveals an intriguing interplay between the human collective above the ice and the fish collective beneath it. Continuous exploitation resulted in lower catch success, despite fish presence increasing over the course of a spot visit. In part, this could be the result of individual differences between fish in catchability. Fish that are more willing to bite (e.g., bolder or larger ones) could be selectively removed, leaving behind shyer individuals that are less likely to bite (48). However, our finding that temporarily leaving a previously exploited area increased the probability of a catch upon return strongly suggests that the fish shoals also encoded recent information on predation risk—a form of collective memory (49). Humans exploited the fact that fish remained present in high numbers at previously successful spots by revisiting these locations later in the competition. Intriguingly, humans may monitor competitors to infer their beliefs and try to preempt their return to previously successful places. Foraging offers an interesting testbed for assessing such theories of mind.

Our work has several limitations. First, the competitive nature of the events may have altered humans’ decision-making strategies relative to humans foraging for subsistence, who rely more heavily on cooperation (50). However, in previous work in our system, incentivizing groups on their collective returns did not fundamentally alter how participants integrated personal and social information when selecting or leaving spots (14), suggesting that our finding are robust to different incentive structures. Second, foragers may have also relied on previous experiences on the lakes, which we did not quantify. Though we intentionally selected lakes that were not well known in the ice-fishing community, future work could study the role of prior information on shaping decision making. Third, because we studied only a single prey species, Eurasian perch, the generalizability of our findings remains unclear. Future studies could expand to other species to investigate whether foragers’ decision making strategies are adaptively tuned to the ecology of different prey species.

To conclude, by tracking human foragers and their prey in the field, our work demonstrates how foragers maximize overlap with a mobile, reactive and invisible prey through the adaptive use of real-time personal and social information, alongside memory. By simultaneously capturing both sides of this interaction, our work empirically reveals how predator and prey collectives co-shape one another’s behaviour —an interplay typically studied theoretically—and illustrates foraging dynamics that likely shaped the cognitive adaptation underlying human hunting throughout evolutionary history.

## Data and code availability statement

All code necessary to reproduce these results is available on GitHub (https://github.com/FelicieDh/FishForagers) and all associated datasets can be dowloaded from Zenodo (https://doi.org/10.5281/zenodo.22798712).

## Acknowledgments

We thank Tomi Keinänen and the North Karelian Sport Fishing Association (Pohjois-Karjalan Urheilukalastajat ry) as well as Petra Siemers-Hering and Katja Münz for their invaluable help organizing the competitions. We acknowledge Julia Chmielnik and Steffi Maukel for their help acquiring the sonar equipment. We are grateful to Veera and Onni from the University of Eastern Finland for help in the field. We further thank Susannah Goss for copy-editing the manuscript and Pietro Nickl for figure-editing.

## Ethics Statement

This study was approved by the Institutional Review Board of the Max Planck Institute for Human Development (A 2024-04). Before participation, each participant signed an informed consent form. We complied with all relevant ethical regulations regarding data protection.

## Authors Contributions

**Conceptualization:** F.D., A.S., V.C., M.M.K., F.F.A.K., D.D., P.R., R.K., P.P., R.H.J.M.K; **Data curation:** F.D.; **Formal analysis:** F.D., A.S., V.C.; **Funding acquisition:** R.K., R.H.J.M.K.; **Investigation:** F.D., A.S., V.C., M.M.K., F.F.A.K., R.K., P.P.; **Methodology:** F.D., A.S., V.C., M.M.K., F.F.A.K., D.D., R.K., P.P., R.H.J.M.K..; **Project administration:** F.D., R.K., R.H.J.M.K.; **Resources:** R.K., R.H.J.M.K.; **Software:** F.D.; **Supervision:** R.K., R.H.J.M.K.; **Validation:** F.D., A.S., R.H.J.M.K.; **Visualization:** F.D.; **Writing - original draft:** F.D., R.H.J.M.K.; **Writing - reviewing and editing:** F.D., A.S., V.C., M.M.K., F.F.A.K., D.D., P.R., R.K., P.P., R.H.J.M.K.

## Methods

### Data collection

We organized 12 ice-fishing competitions over six days (one in the morning and one in the afternoon) between 15 and 22 March 2024 in Northern Karelia, eastern Finland. Competitions were held on three lakes: Sompalampi (62.6209°N, 29.5248°E, 15 and 18 March), Kuorinka (62.5935°N, 29.3937°E, 19 and 20 March), and Pitkänen (62.6511°N, 29.5643°E, 21 and 22 March) (Extended Data Fig. 1). These lakes were used as they were relatively unfamiliar to participants, predominantly contained Eurasian perch, and ranged mainly between 4m and 15m in depth. Initial sonar testing indicated that this depth range was most suitable for fish detection.

At each lake, we created four non-overlapping arenas, each of which was used once. Each arena consisted of three overlapping circles with a diameter of 58m, with all three circles intersecting at one central point (Fig. 1C) (arena size approx. 7,000m^2^). A sonar was placed at the center of each circle to provide complete coverage of fish presence in the arena (see below).

In total, 18 unique participants took part in this study (15 men, 3 women; mean age = 69, min = 56, max = 80). Each competition involved six participants, and every participant competed in four competitions: two on a single day (morning and afternoon) and two more, two days later. This design ensured that each participant fished on two different lakes and in four distinct arenas. **RK** recruited the participants, only inviting highly experienced anglers who regularly participated in ice-fishing competitions (mean number of years of ice-fishing experience = 53 (SD = 8); mean number of self-reported ice-fishing competitions per year = 36 (SD = 23)).

Each competition lasted two hours (from 10:00 to 12:00 or 13:00 to 15:00), during which participants were free to fish and relocate within the arena. Participants used their own rods and lures. We only authorized manual drills with a diameter up to 155mm (i.e., no power drills) to standardize the effort needed to create new fishing holes. Only fly larvae (*Calliphora spp.*), bloodworms (*Chironomus spp.*) and redworms (e.g., *Eisenia fetida*) were allowed as bait; all bait had to be firmly attached to the hook (i.e., using loose bait to attract fish was not allowed). Participants were instructed to target Eurasian perch (*Perca fluviatilis*) and to release any other species caught. Only one fish of a different species (*Esox lucius*) was caught during the competitions.

### Incentivization

Participants were reimbursed based on total catch weight (of Eurasian perch to the nearest gram) in line with local official competitions. Monetary prizes were as follows: €60 for first place, €40 for second place and €20 for third place. The maximum amount a participant could earn in a single day was thus €120. The total awarded over the six days was €1,440.

### Human foraging behaviour

Each participant was equipped with a sports watch (Garmin Forerunner 245) that continuously recorded their spatial positioning at a sampling frequency of 1Hz (using the “run” mode) and with a head-mounted camera (Drift Ghost XL) positioned to capture their fish catches. To synchronize data streams, we filmed the time on the watch for ∼10s before each competition. Horn signals marked the start and end of each competition, providing an additional audible timestamp in the video recordings. Cameras and watches were collected and their data downloaded every evening.

We manually annotated each video using the BORIS software (1), annotating three behavioural states (relocating, drilling and angling) and one event type (fish catch). Full details on video annotation can be found in (2). The extracted behavioural states and fish catch events were merged with the timestamped GPS positioning data to produce a high resolution spatiotemporal behavioural dataset for each participant and each competition (Fig. 1C).

### Fish presence estimations

We used three sonar transducers (Lowrance ActiveTarget2) connected to 12” monitors (Lowrance HDS pro) via a module (Lowrance ActiveTarget2) to track fish in the arenas. The monitors were mounted on the lid of a 25l polystyrene cooler. The module and a portable battery (40Ah, 12.8V, Lithium LiFePO4) powering the devices were placed inside the cooler, insulating them from the cold. Sonar setups were transported across the lake surface by sled. Transducers were mounted in forward mode on a custom ice-fishing mounting tripod supporting a rotating shaft that could be lowered up to 1m beneath the ice surface, allowing full 360° rotation. An orange tab, lined up with the transducer, was affixed to the top of the rotating shaft to track the transducer’s orientation beneath the ice. A magnetic compass, calibrated for northern latitudes, was attached to the orange tab to take bearing measurements. A small tent shielded the monitors from sunlight and harsh weather and prevented participants from seeing the screens when fishing nearby.

Each sonar was used to visualize the lake bottom and the fish present in the water column up to a distance of ∼35m and at a beam angle of 18°. The design of the arenas ensured complete sonar coverage (Fig. 1C). The sonar units were installed in an equilateral triangle with 50m sides approx. one hour before each competition, using three 50m paracords. We drilled holes through the ice at the three corners of the triangle (i.e., at the center of each circle) using a 255mm cordless power auger, and installed tripods such that the transducers were ∼20cm under the ice (average ice thickness = 55cm (SD = 3.1)). We then checked whether the sonars were working properly and that there were no obstructions (e.g., underwear debris) present that would obscure fish detections. If obstructions were present, the sonars were moved until an appropriate placement was found. We used a handheld GPS to record the exact coordinates of each sonar. We drilled a small hole ∼2m away from the sonar in the direction of the geomagnetic North (0°) and lowered a metal nut attached to a rope ∼50cm below the ice. The nut was visible on the sonar’s monitor and was used to determine when a full rotation of the transducer was completed. The arena limits were then drawn in a 29m radius circle around each sonar (using a paracord). The edges of the arenas were marked by small colourful cones every 10 m.

Each sonar was operated by two observers: one controlled the transducer and the other noted down the fish observations following a standardized protocol. The orange tab (and by extension the underwater transducer) was first oriented North (as indicated by the metal nut on screen and 0° on the compass) and then slowly rotated clockwise until fish were detected on the display. When fish were detected, the observer recorded the time, transducer bearing, distance from the transducer to the fish (for a fish shoal, min and max distance), depth of the fish (for a fish shoal, min and max depth) and estimated the number of fish. Distance and depth were obtained by clicking on the fish on the monitor. Fish were counted individually up to 20; estimates between 20 and 100 were rounded to the nearest five and estimates greater than 100 were rounded to the nearest 50. Once the transducer faced North again, we rotated it 360° counterclockwise and started a new scanning rotation. We recorded the time the transducer faced North at the start of each scanning rotation to quantify rotation duration. Sonar displays were mirrored to Apple iPad Pro devices set to screen recording, creating video copies.

For each fish detection, we used the transducer’s GPS coordinates, the transducer’s bearing, and the average distance of the fish to the transducer to calculate the GPS coordinates of the centre of the fish shoal (or individual fish). Fish shoal radii were calculated as half the difference between the maximum and minimum distances from the transducer, assuming a circular horizontal shape. Based on the estimated shoal size, we then randomly generated the corresponding number of fish positions within this radius. This resulted in a uniform distribution of fish around the centroid, providing a single coordinate for each fish at each time point. The resulting dataset thus contained easting, northing, and time for each individual fish detected.

### Fish presence modelling

The dataset from the sonar provided instantaneous fish detections during each rotation, but not continuous fish presence data for the entire arena (as the transducer had a horizontal beam of 18°). To represent fish presence continuously in space and time, we used GP models (3). Each fish coordinate served as a spatiotemporal observation **x***_i_* = (*E_i_, N_i_, t_i_*), and the GP applied a kernel that decayed with increasing spatial and temporal distance from that observation. This approach generated a dynamic intensity surface in which areas close in space and time to a fish detection retained high values, while locations further from previous detections showed progressively lower values.

GP models were performed individually for each competition using the scikit-learn library (4) in Python 3.11 (5). The covariance structure of the GP was defined as the product of a constant kernel, *C*, and a radial basis function (RBF) kernel:

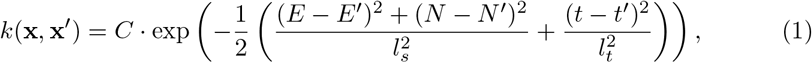

where *l_s_* is the identical spatial length scale applied to both easting and northing (in metres) and *l_t_* is the temporal length scale (in minutes). This enabled the model to capture both global scaling effects and smooth variations along spatial and temporal dimensions. The constant kernel was initialized with a value of 1.0, and the RBF kernel with length scales of 1.0 for each input dimension.

We first optimized hyperparameters separately for each competition to account for competition variability in fish presence. Input features were easting, northing, and time since competition start (in minutes); the target variable was set as an array of ones (**y** = **1**), representing confirmed fish presence at each detected coordinate. Under this formulation, the GP does not perform regression in the conventional sense; instead, it interpolates a continuous spatiotemporal fish presence surface from the discrete detection events.

The data for each competition were split into training (80%) and test (20%) subsets without shuffling, preserving temporal order. Hyperparameters (constant kernel value, RBF length scales along each dimension, and the noise term *α*) were optimized in two stages. First, a coarse randomized search explored broad parameter ranges. For the spatial parameters, the lower bound was taken as 2.3 m, reflecting the sonar’s angular resolution: With an 18° beam, an object at a 15m range would first enter the beam (and therefore be detectable) roughly 2.3m before the beam was pointed directly at it. The upper bound of 30m represented the maximum distance from a participant to the sonar. For the temporal parameters, we took the 1^st^ quartile of the scan lengths (1.8 min) and the maximum scan length (i.e., time to complete a sonar rotation, 46.5min). Second, a finer search was conducted over a narrowed parameter space informed by the coarse results. In both stages, we used randomized search cross-validation with 30 iterations. Time series cross-validation was implemented using four sequential splits to ensure that training data always preceded test data, preventing information leakage from future observations. Model performance during tuning was evaluated using negative mean squared error (MSE). The best hyperparameters for each competition were recorded, and the corresponding best estimator was refitted on the full dataset for that competition, including previously held-out test data. Predictions on the held-out test subset were used to compute MSE, providing a measure of model accuracy.

After hyperparameter optimization, competition-specific GP models were refitted using the full data and the previously optimized parameters (Supplementary Table 14). The kernel was reconstructed using the optimized constant value and RBF length scales, with bounds fixed to prevent further optimization. The noise term (*α*) was set according to the optimized value.

Following model fitting, we generated spatial-temporal predictions of fish presence for each competition. For each competition, the GP model was loaded and a uniform 2D spatial grid with 1m² grid cells and 135 sampling points along each axis was defined. Predictions were performed for each grid cell and for each minute of the competition (defined as a new spatiotemporal point **x***^∗^*) using the GP posterior predictive mean:

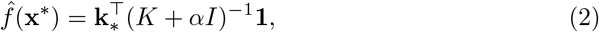

where **k***_∗_* is the covariance vector between **x***^∗^* and the training data, *K* is the training data covariance matrix, and **1** is the target vector of ones representing all confirmed fish detections. The obtained value *f̂*(**x***^∗^*), ranging from 0 (i.e., highly certain no fish is present) to 1.2 (i.e., highly certain a fish is present) is referred to as ‘fish presence’.

### Angling strategy validity

To understand if the spatiotemporal predictions generated by the GP models predicted individuals’ foraging success, we tested whether successful spots (i.e., at least one fish catch) had higher fish presence at spot arrival than unsuccessful spots (i.e., no fish catch). We fitted a linear regression with fish presence at spot arrival as response variable and success (0,1) and distance to the sonar (m) as population-level effects, with a random intercept for each competition. To test if active angling locations had higher fish presence than average in the arena, we estimated the average fish presence per minute across the entire arena and the average fish presence within 3m of each participant angling in that minute. We then fitted a linear regression with average fish presence as response variable and minute and location (categorical: whole arena or within 3m of participants) as population-level effects, adding a random intercept for each competition and a random slope over minutes. Estimates for this analysis are presented in Extended Data Fig. 3B.

All linear models were built in *brms* (6) with default priors. All models were run for 2000 sampling iterations following 1000 warmup iterations.

### Spot selection model

To identify the spatial features driving spot selection decisions, we extended the computational modeling approach introduced by (2). We modeled each spot choice as a discrete selection among a set of candidate spots. For each observed spot choice, we constructed a choice set consisting of the actual chosen spot alongside *S^∗^ −* 1 = 30 counterfactual alternatives. Counterfactual locations were generated by sampling step lengths and turning angles from the empirical distributions, with draws constrained such that the resulting positions fell within the boundaries of the arena.. This approach, analogous to step-selection functions used in movement ecology (7), ensures that simulated alternatives reflect realistic movement constraints.

### Choice model

Let *t ∈ {*1*,…, T}* index successive spot choices within a foraging trip consisting of *T ∈* N visited angling spots. For each angling spot *t*, let *S*_[*t*]_ *∈ {*1*,…, S^∗^}* index chosen locations within each choice set *S^∗^ ∈* N. We modeled the probability of selecting the observed spot as:

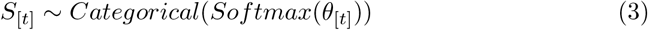

where *θ ∈* R*^S∗^* is a vector of length *S^∗^*and the value assigned to each candidate spot *s* at time *t* is a linear combination of spatial features:

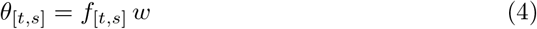

Here, **w** *∈* R*^M×^*^1^ is a vector of *M* feature weights to be estimated, and *f*_[*t,s*]_ *∈* R^1^*^×M^* is a row-vector of spatial features evaluated at candidate spot *s* at the time of spot choice *t*. A positive weight indicates that the participant selects locations with higher values of the corresponding feature relative to the other available locations; a negative weight indicates avoidance.

### Spatial features

We computed three kernel-based features capturing different sources of information, and three additional features to control for relevant movement and ecological context, yielding *M* = 6 features in total.

### Social information

The social feature quantifies the local density of other participants near each candidate spot at the moment of spot choice. For each candidate spot *s*, this is computed as a sum of Gaussian kernel weights over the *N*_[*t,s,*1]_ observed competitor positions:

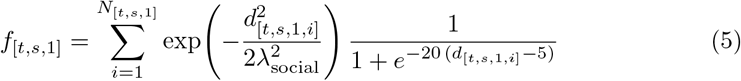

where *d*_[*t,s,*1*,i*]_ is the Euclidean distance from candidate location *s* to the *i*-th competitor at time *t* corresponding to the start of angling, and *λ*_social_ is a spatial bandwidth controlling how rapidly social influence attenuates with spatial distance. The logistic term smoothly suppresses the contribution of competitors closer than 5m, reflecting the minimum–distance rule between participants enforced in ice-fishing competitions.

### Personal success and loss

The success and loss features capture proximity to locations where the focal participant has previously caught fish (success) or failed to do so (loss). For these personal information features, we incorporated both a spatial (in metres) and a temporal (in number of spots sampled since) decay function, using a product of Gaussian kernels:

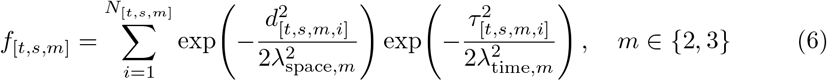

where *d*_[*t,s,m,i*]_ and *τ*_[*t,s,m,i*]_ are the spatial and temporal distances from candidate location *s* to the *i*-th prior event of type *m* (success = 2 or loss = 3), and *λ*_space*,m*_ and *λ*_time*,m*_ are the corresponding bandwidths. When no prior events of a given type were available, the corresponding feature was set to zero. Success and loss were estimated with separate spatial and temporal bandwidths.

### Distance to the sonar

Because participants preferred to remain at a distance to the sonar during competitions, we controlled for the Euclidean distance from each candidate location to the closest sonar, 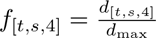, with *d_max_* being the maximum distance from the sonar, and smaller values indicating locations closer to the sonar (0 to 1).

### Locality features

To control for individual and lake-level differences in relocation distances, we included *f*_[*t,s,*5]_ = *r*_[*t,s*]_, where *r*_[*t,s*]_ is the step length to candidate location *s*. To capture potential changes in movement behaviour over the course of the competition independently of information use, we included an interaction term *f*_[*t,s,*6]_ = *f*_[*t,s,*5]_ *· j*_[*t*_*_−_*_1]_, where *j*_[*t*_*_−_*_1]_ *∈* [0, 1] is the elapsed competition time upon arrival at a spot, normalized by the total competition duration.

### Prior specification

All five bandwidth parameters were estimated on the log scale:

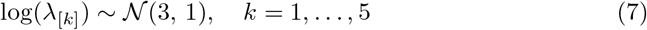

Feature weights were assigned weakly informative priors:

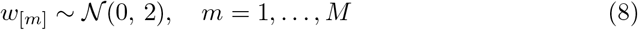

### Hierarchical structure

To account for the nested structure of the data, we estimated all feature weights with random offsets at three grouping levels: competition, individual and foraging trip (i.e., an individual in a competition). Bandwidth parameters received only competition-level offsets. At each grouping level, offsets across all parameters within a group were drawn jointly from a multivariate normal distribution:

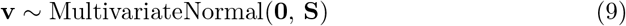

where the variance-covariance matrix is decomposed as:

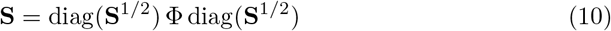

and Φ is the correlation matrix among parameters. To improve sampling efficiency and avoid funnel-shaped posterior geometries, we used a non-centred parameterization (8) via Cholesky decomposition. Letting *L* denote the Cholesky factor of Φ such that Φ = *LL^T^*, and *z ∼* MultivariateNormal(0*, I*), the group-level offsets are estimated as:

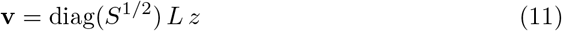

We placed an LKJ prior on the Cholesky factor, *L ∼* LKJ(4), which regularizes extreme correlations among parameters (9). SDs were assigned exponential priors, 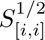 *∼* Exponential(1).

### Computation

The model was implemented in Stan (10) and fitted via Hamiltonian Monte Carlo using CmdStanR (11). Within-chain parallelization of the categorical log-likelihood was achieved using Stan’s reduce sum function. Convergence was assessed using the Gelman-Rubin statistic, with *R̂ ≤* 1.01 required for all parameters (12; 13). Posterior estimates are presented in Supplementary Table 2.

### Ecological validation of spot selection

To examine how well the spatial features predicted fish presence, we used a Beta regression to model the fish presence at the chosen location *S*_[*t*]_. At each spot chosen by a participant, we computed the social, success and loss features using the mean bandwidths estimated by the spot selection model for the corresponding competition. We further extracted the distance to the sonar, time in the competition and average fish presence in a radius of 3m around the spot upon participant’s arrival. We then modelled fish presence (scaled to fit a beta distribution) as a function of the social, success and loss features, distance to the sonar and time in the competition with a global random intercept per competition and a random slope on time. Distance to the sonar was included as a control as fish are easier to detect closer to the sonar. All explanatory variables in the model were standardized at the competition level to have a mean of 0 and an SD of 1. Posterior estimates are presented in Supplementary Table 3.

### Spot leaving model

To examine how participants decide to leave their current fishing spot, we implemented a hierarchical Bayesian logistic regression model predicting the binary outcome of staying (0) or leaving (1) for each 10-second time bin spent at a spot as a function of a set of spatial and temporal features. The model was computationally implemented similarly to the spot selection model above.

### Model structure

The leaving decision at each time bin *t* within a spot visit was modeled as a Bernoulli trial:

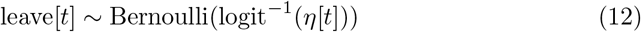

where *η*[*t*] is a linear predictor. We estimated two versions of *η*[*t*] depending on whether at least one fish had been caught at the current spot:

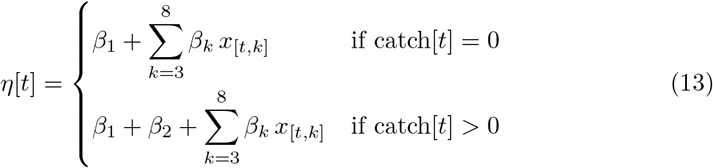

where *β*_1_ is a baseline leaving probability (intercept), *β*_2_ captures the shift in leaving probability associated with having caught at least one fish at the current spot by time *t* (fish catch), and *β*_3_*,…, β*_8_ are weights for the remaining predictors *x*_[*t,*3]_*,…, x*_[*t,*8]_ described below.

### Predictors

The predictor vector *x*_[*t*]_ contained the following variables: *x*_[*t,*3]_, local fish presence at the current spot, computed as the average fish presence in a 3m radius around an active fishing spot; *x*_[*t,*4]_, time since the last event (fish capture or arrival at the spot, in seconds), describing the effect of time without a catch; *x*_[*t,*5]_, distance to the nearest sonar (0,1), included as a control variable because participants tended to remain near the arena boundary during competitions; *x*_[*t,*6]_, social density feature, representing the density of competitors; *x*_[*t,*7]_, personal success feature, reflecting proximity to previously successful locations; *x*_[*t,*8]_, personal loss feature, reflecting proximity to previously unsuccessful locations. For *x*_6_, *x*_7_, and *x*_8_, features were computed in advance using the mean posterior bandwidth parameters estimated in the spot selection model.

### Sequential model comparison

To assess the relative contribution of each set of predictors to participants’ ability to track fish presence, we sequentially removed predictors from the full model: first social information; then success and loss information; and finally catch information. If removing a predictor increased the fish presence coefficient, this indicated that the removed predictor had independently encoded information about fish presence in the full model (Extended Data Fig. 5B).

### Prior specification and computation

Feature coefficients were assigned weakly informative priors *β_k_ ∼ N* (0, 1). The hierarchical random effects followed the same non-centred parameterisation, LKJ correlation priors, and exponential scale hyperpriors as described for the spot selection model, with offsets estimated at the competition, individual, and trip level. The model was fitted in Stan (10) via CmdStanR (11), with within-chain parallelization implemented using reduce sum on foraging trips. Convergence was confirmed using *R̂ ≤* 1.01 for all parameters. Posterior estimates are presented in Supplementary Table 4.

### Ecological validation of spot leaving model

Similar to the spot selection model, we examined how well the spatial and temporal features predicted fish presence, including all features described above except fish presence, which became our response variable. Because fish presence was calculated per minute, we used 1 min time bins. For each minute spent at a spot we modeled the fish presence (scaled to fit a beta distribution) as a function of the social, success and loss features, distance to the nearest sonar, fish catch (0,1), and time since the last event (i.e, fish catch or arrival at the spot, in minutes) with a random intercept per competition. As in the validation of the spot selection model, we used the spatial feature’s mean bandwidths. The three features and distance to the sonar were then standardized at the competition level to have a mean of 0 (SD = 1). Posterior estimates are presented in Supplementary Table 5.

### Effect of angler presence on fish presence

To quantify the effect of participants above the ice on fish presence beneath it, we compared the dynamics of fish presence between visited spots and control spots. We used the following procedure to identify control spots. For each arena, we created a grid of 100 locations and calculated for each location and each minute (i) average fish presence and (ii) whether any participants were present within a 10m radius. This radius exceeds the spatial length scale (5.3–7.1m) estimated by the GP, meaning that fish presence at these control locations is expected to show substantially reduced spatial correlation with nearby angler positions, approximating natural (undisturbed) fish behaviour. At the start of each angling event (i.e., a participant’s arrival at a new spot), we identified a set of candidate locations meeting the following criteria: (1) Its fish presence at the corresponding time point was within 0.02 of the fish presence at the location of the angling event (at the start of angling). This threshold, corresponding to ∼2% of the full range of the fish presence was chosen to ensure close correspondence in initial conditions between visited and control spots while retaining a sufficient number of candidate matches per angling event. (2) It occurred at the same time (i.e., within the same minute of angler arrival) and in the same arena as the angling event. (3) No participant was present within a radius of 10m at any point during the angling event. From this set of candidate locations, we randomly selected up to five as control spots for each angling event (mean = 3.6, median = 4, min = 1, max = 5, depending on availability).

We then investigated the temporal change of fish presence at visited spots relative to control spots, separately for successful and unsuccessful spots. This separation was motivated by two factors: the effect of participants on fish presence may differ by outcome (e.g., fish removal may directly reduce local density or change fish behavior) and visit duration was significantly shorter at unsuccessful than at successful spots. For all visited and control spots, we calculated Δfish: the difference in fish presence relative to the start of the visit (first minute). Positive (negative) values indicate that fish presence increases (decreases) over time. We then modeled how fish presence developed at (un)successful spots relative to control spots (controlling for spot visit duration). In both cases, we modeled Δfish as a function of a three-way interaction between time (minutes), fish presence at minute 1, and spot type (“visited” vs. “control”). We also included distance to the sonar to control for potential detection effects and used random intercepts for each competition and each spot. Posterior estimates are reported in Supplementary Tables 6, 7; a sensitivity analysis for control spot selection is shown in Extended Data Fig. 6A, B.

### Effect of angler presence on catch probability

To test how catch probability changed over successful spot visits, we recorded fish presence upon arrival and whether a fish was caught in each subsequent minute. We then modeled how fish presence and time since visit start affected catch probability, controlling for fish presence at spot arrival and distance to the sonar, using a Bernoulli regression with random intercepts for competition and spot. Posterior estimates are presented in Supplementary Table 8.

### Catch recovery in the absence of anglers

To test whether catch probability recovers when an area is not fished for a period of time, we split each arena into a 15m Voronoi grid and computed for each cell and minute whether a participant was present and the number of participants present. Consecutive minutes with at least one participant in a cell were grouped into discrete visit events. When a cell was revisited after a gap, the event was assigned an ordinal index reflecting its position in the sequence of visits to that cell (i.e., first visit, second visit, etc.). For each event, we derived the following variables: whether any participant caught a fish during the event (binary, 0/1); time away, defined as the duration between the end of the preceding event and the start of the current event in the same cell (in minutes), set to 180 minutes for the first visit to a cell (corresponding to the minimum time a spot received no visit, given our presence on the lake setting up the experiment); time elapsed within the current event (in minutes); the maximum number of participants to have fished in the cell during the event; and mean fish presence, calculated as the average fish presence in the cell across the event duration.

We then modeled catch success (binary, 0,1) for each event and minute as a function of time in the event, event number (first, second, etc. visit), time away, number of participants and average fish presence, with a random intercept for cell identity. Posterior estimates are presented in Supplementary Table 9 and visualized in Extended Data Fig. 7; a sensitivity analysis on grid size is reported in Extended Data Fig. 6C.

### Revisit analysis

We used a clustering algorithm to test what triggered participants to return to previously visited spots. We used DBSCAN (14) with an *ɛ* of 10m (diameter from the spot) to identify clusters of spots (Fig. 5A). Spots within 10m of a previously visited spot were assigned to the same cluster. Spots more than 10m away from any previously visited spot were assigned a new cluster. Clusters were updated after each spot departure.

To determine how features of a spot visit impact a participant’s likelihood of returning to the same cluster, we calculated, for each spot departure, whether the participant was successful or not during that visit, and mean fish presence within 3m of the spot at the time of departure. Continued exploration within the same cluster did not qualify as departure (Fig. 5A). For each departure, we coded a binary outcome indicating whether the participant subsequently revisited the cluster (1) or not (0); multiple revisits after a given departure were coded as a single event rather than a count. We then modeled likelihood of returning to the cluster that a spot belongs to as a function of visit success, and ran a separate model testing return likelihood as a function of fish presence at departure. Both models included the minute of departure (1–120) to account for temporal changes in exploration tendency, and a random intercept for each competition. Posterior estimates are presented in Supplementary Tables 10, 11.

To determine how recent success or loss affected the likelihood of revisiting a previous cluster, we determined for each spot visit whether it was successful or not and the fish presence upon departure. We then determined whether the next spot visited by a participant was a revisit to a previously visited cluster. We modeled the likelihood that the next spot visit would be a cluster revisit as a function of visit success, and ran a separate model testing this likelihood as a function of fish presence at the last spot visit. Both models included the time at which the spot choice was made and the number of spots available for revisits in the arena as predictors, with a random intercept for each competition. Posterior estimates are presented in Supplementary Tables 12, 13; a sensitivity analysis on *ɛ* values is presented in Extended Data Fig. 6D–G.

## Extended data

**Extended Data Figure 1.**
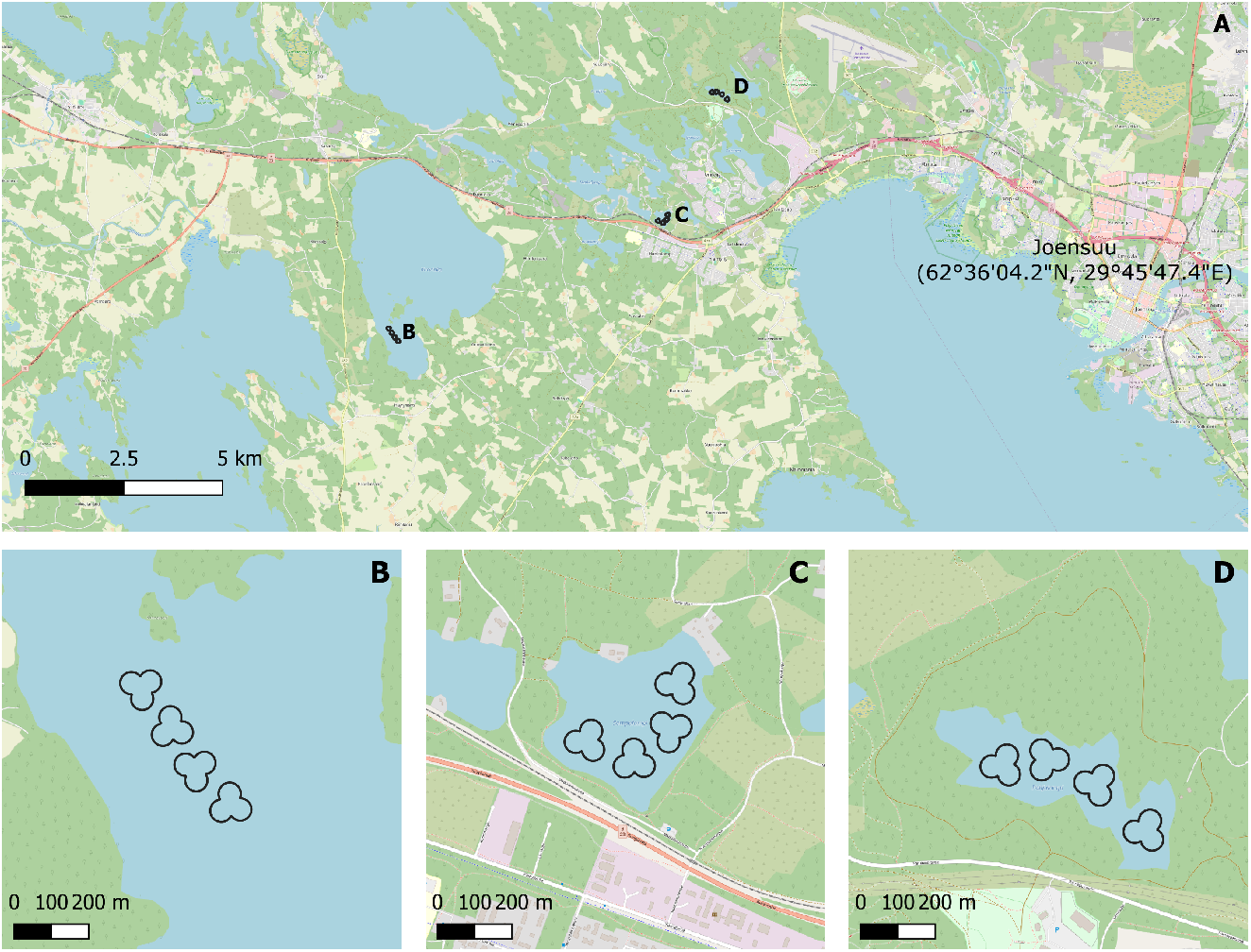
Study area. (A) Map of the study area, showing the city of Joensuu (Finland) in the East, and the three lakes used for the experiment: (B) Lake Kuorinka. (C) Lake Sompalampi. (D) Lake Pitkänen. (B–D) show the position of the four arenas on each lake.

**Extended Data Figure 2.**
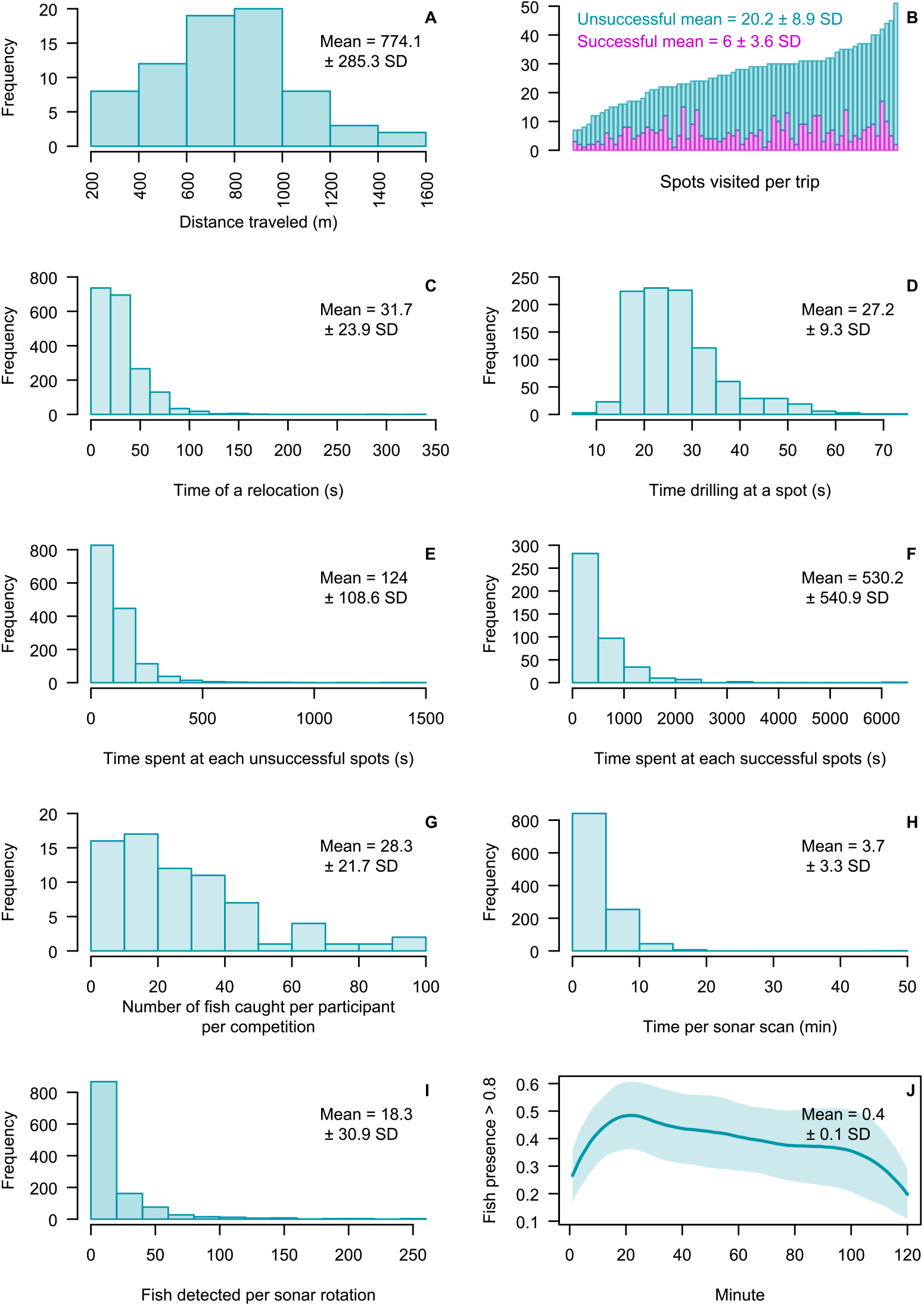
Data description. (A) Distribution of distances traveled per participant across all competitions. (B) Number of successful (i.e., at least one fish caught) and unsuccessful (i.e., no fish caught) angling spots visited per trip (i.e., per participant per competition). Overall, participants visited 26.3 *±* 9.3 spots per trip. (C) Distribution of the time spent per relocation per participant across all competitions. (D) Distribution of the time spent drilling per participant across all competitions. (E) Distribution of the time spent fishing at unsuccessful spots per participant across all competitions. (F) Distribution of the time spent fishing at successful spots per participant across all competitions. (G) Distribution of the number of fish caught per participant across all competitions. (H) Distribution of the time spent per sonar rotation across all competitions. (I) Distribution of the number of fish detected per sonar rotation across all competitions. (J) Proportion of the arena where fish presence as estimated by GP models was above 0.8 over time across all competition. The line represents the average fish presence per minute; the ribbons show the SD.

**Extended Data Figure 3.**
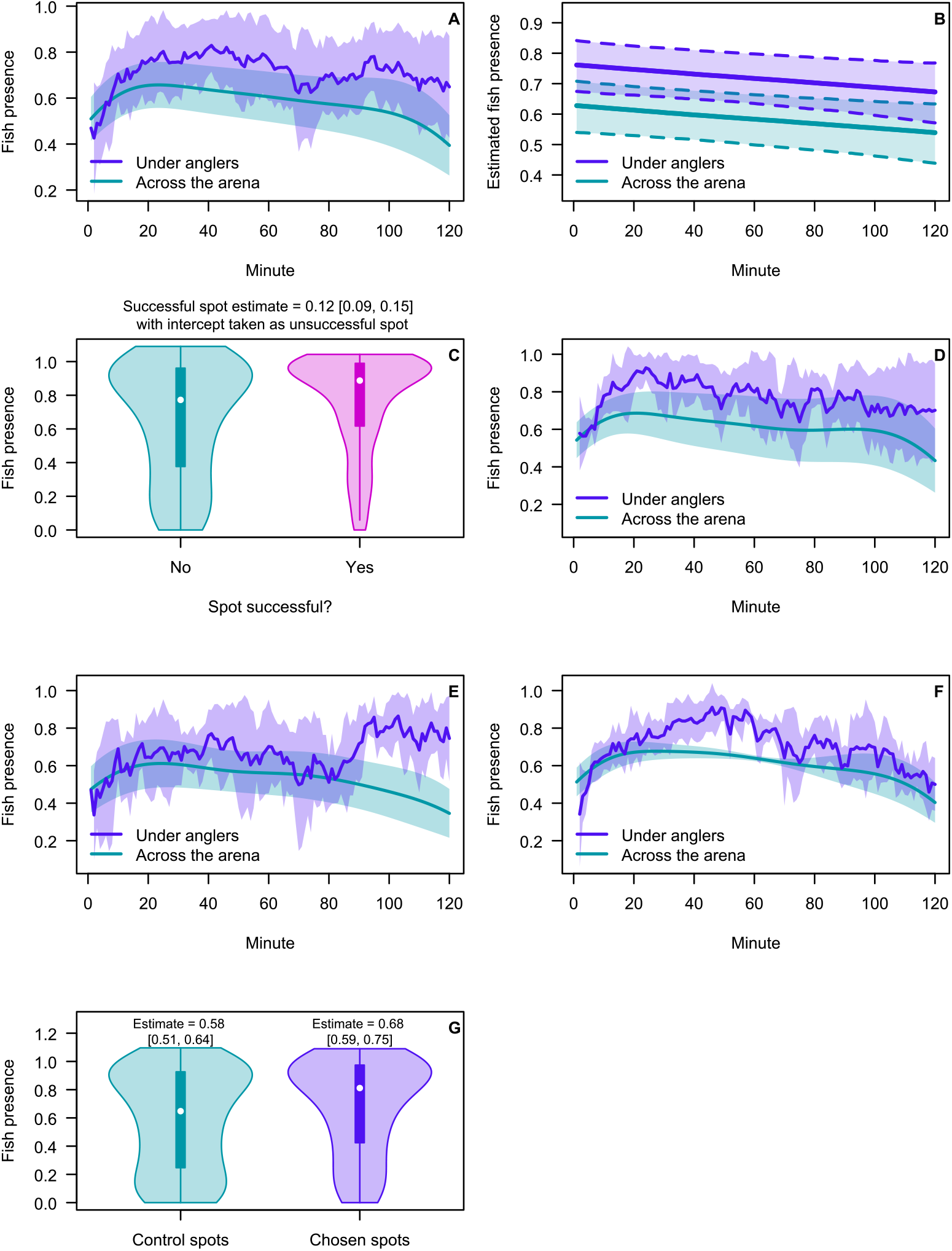
Fish presence description. (A) Mean fish presence across the whole arena (mean *±* SD) and at active angling spots overall. (B) Estimates of fish presence in the arena and at active angling spots. These results were obtained by modeling fish presence as a function of location (under anglers or not) and minute, with a random intercept for each competition and a random slope over minutes. Fish presence was lower where there were no anglers (Estimate [95% credible interval (CrI)]=*−*0.13[*−*0.14*, −*0.13]) and decreased over time (Estimate [95% CrI]=*−*0.0007[*−*0.001*, −*0.00002]). (C) Mean fish presence at successful and unsuccessful spots. Fish presence was measured at the moment of a participant’s arrival. White dots represent the median; the box extends from the first to the third quartile. (D) Mean fish presence across the whole arena (mean *±* SD) and at active angling spots for Lake Kuorinka; (E) for Lake Sompalampi; (F) for Lake Pitkänen. (G) Fish presence at spots chosen by participants (measured at the moment of a participant’s arrival) and at control spots. We randomly generated five control spots per chosen spot. Estimates calculated using a linear regression with fish presence as response variable, spot type (control or chosen) as predictors, and a random intercept per spot choice.

**Extended Data Figure 4.**
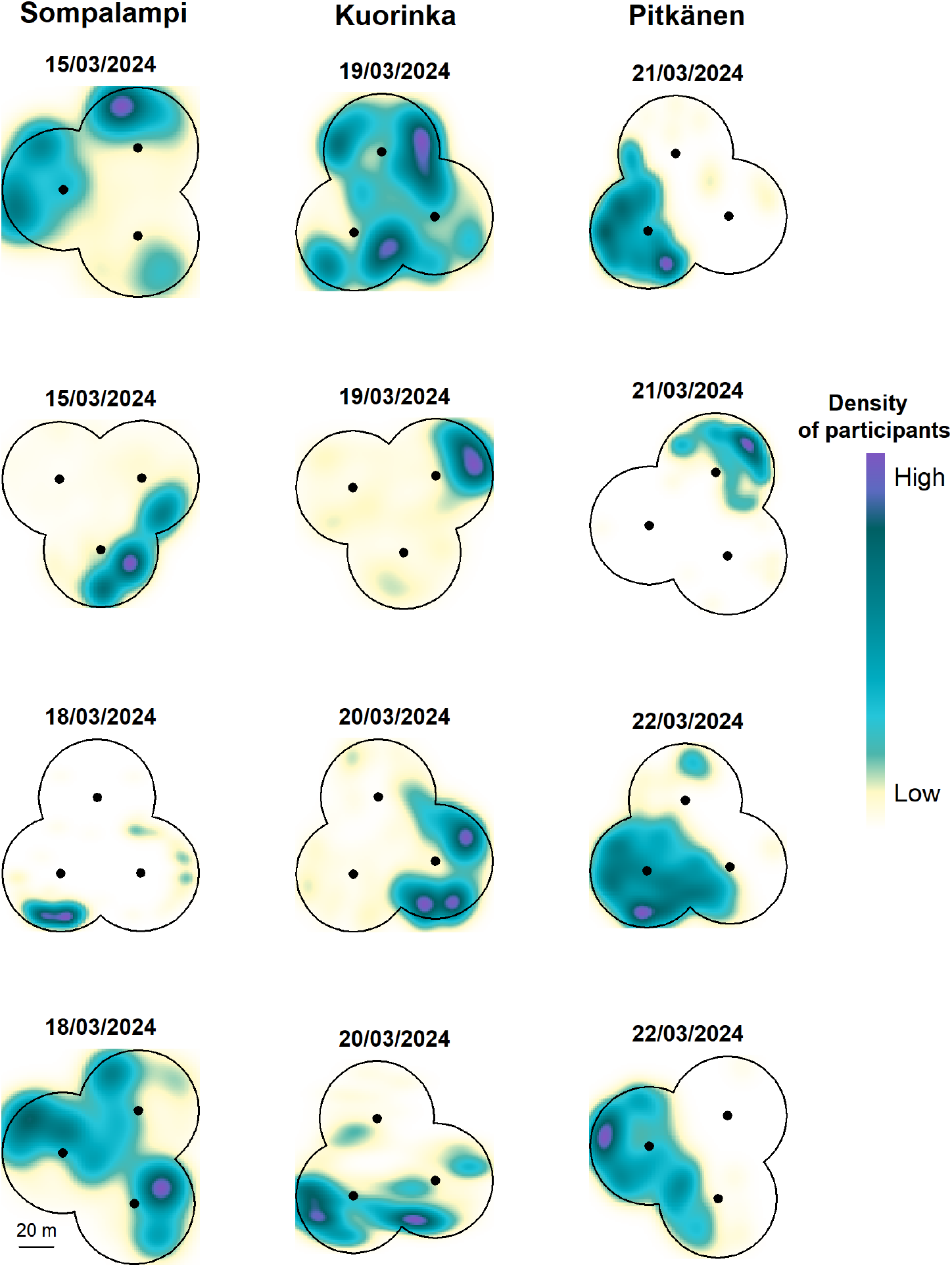
Density heatmaps of area use for each competition. The density was estimated using kernel density estimations in the MASS package (1), using one data point per angling minute per participant. Black dots show sonar locations.

**Extended Data Figure 5.**
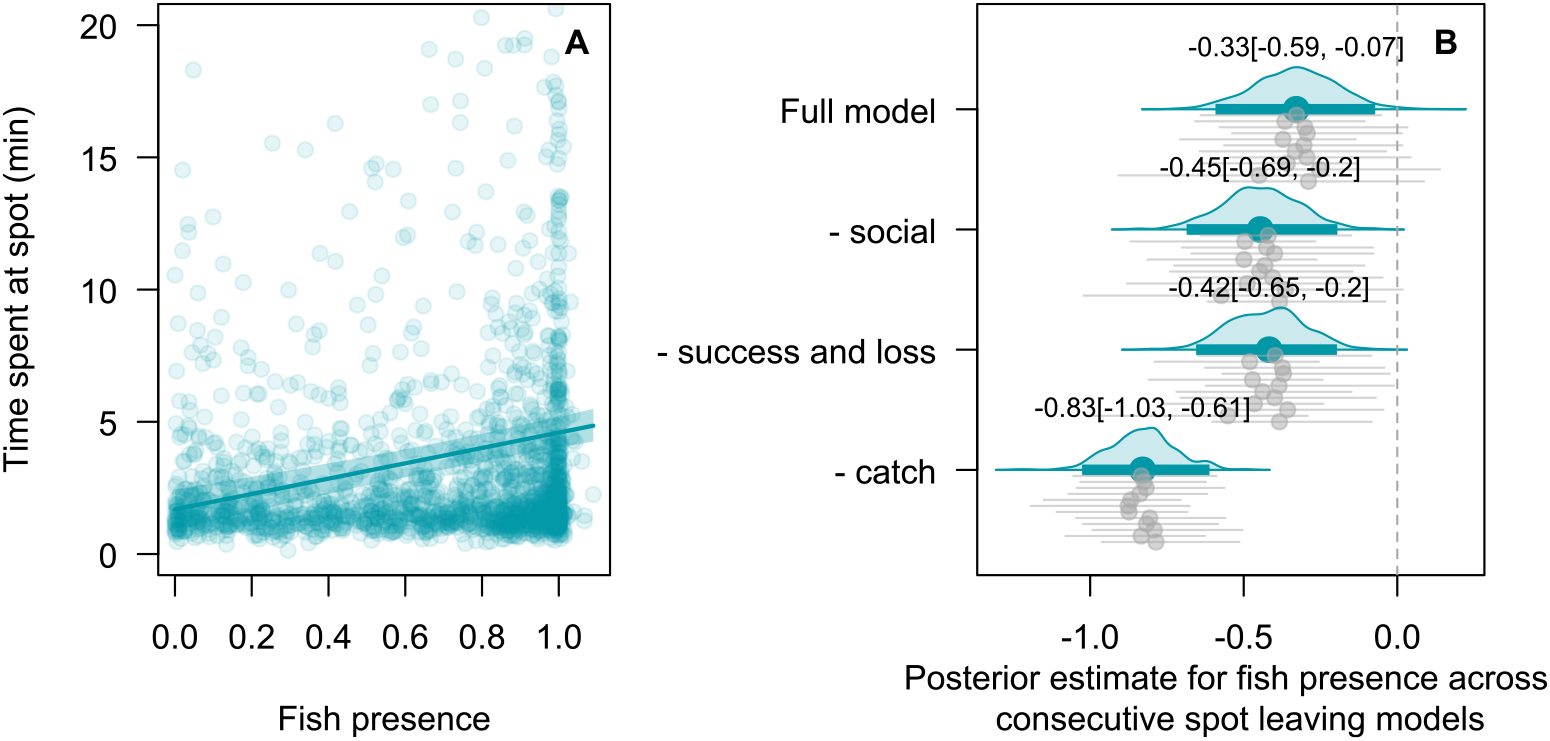
Spot leaving. (A) Time spent at a spot as a function of fish presence at arrival. Participants spent more time at spots with greater fish presence (Estimate [95% credible interval]=2.9[2.06, 3.72]). The estimate derives from a linear regression with time spent at a spot as response variable, fish presence and distance to the nearest sonar as a predictors, and a random intercept per competition. (B) Effect of fish presence on spot leaving decisions in consecutive models. The full model is presented in the main text. In subsequent models, we first removed the social feature, then the success and loss feature and finally the fish catch information (i.e., catch, yes or no), leaving only distance to the sonar and fish presence as predictors. Removing predictors leads to an increasingly strong negative effect of fish density on leaving likelihood, suggesting that participants use these predictors to infer fish presence. Estimates and 95% credible intervals are reported on the figure.

**Extended Data Figure 6.**
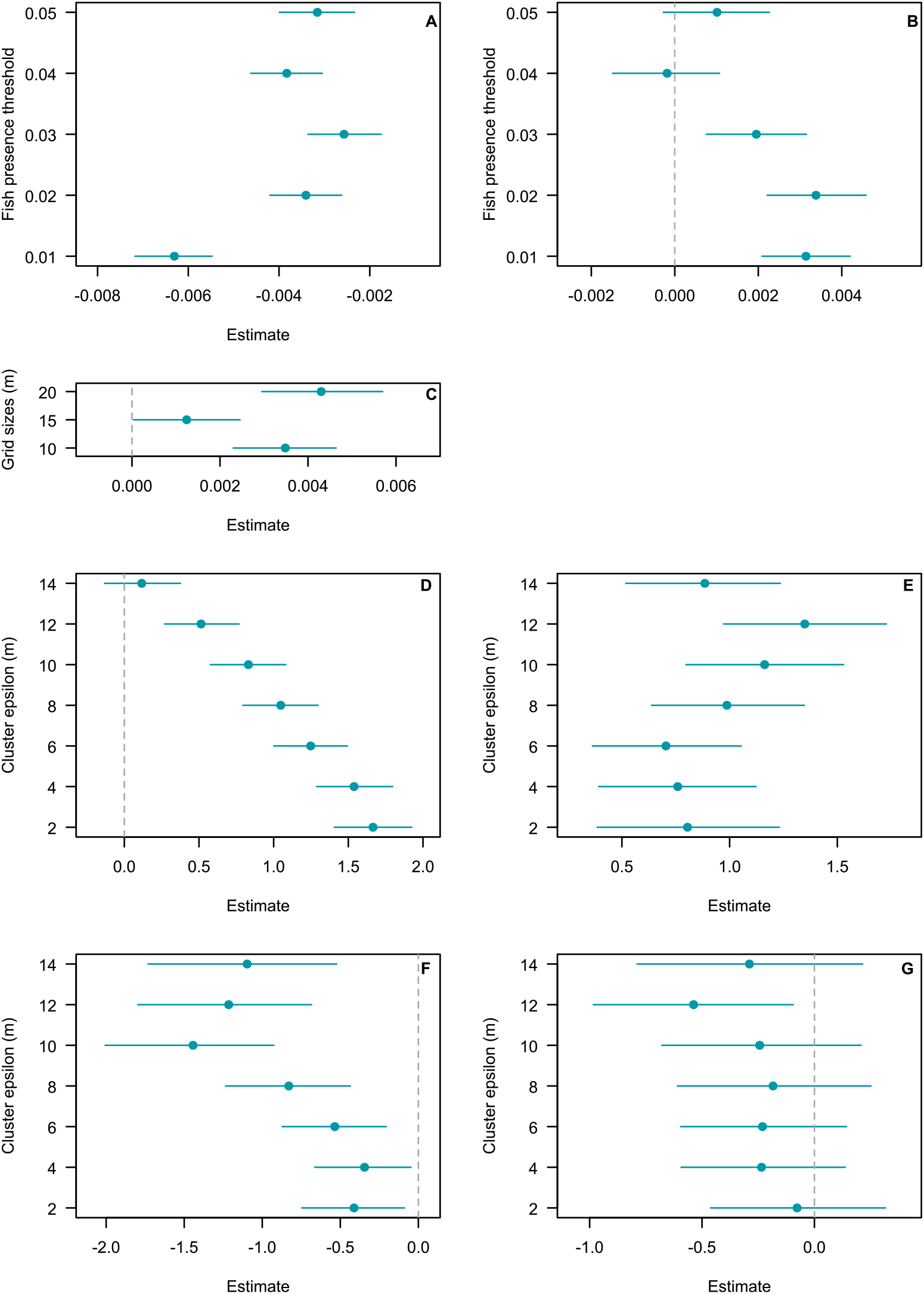
Sensitivity analyses results. (A) Predicted change in fish presence over time at unsuccessful spots across different fish presence thresholds. (B) Predicted change in fish presence over time at successful spots across different fish presence thresholds. (C) Effect of time spent away from an area on catch recovery across different grid sizes. (D) Effect of success (catch) at a spot on the probability of it being revisited across different *ɛ* clustering thresholds. (E) Effect of fish presence when leaving a spot on the probability of it being revisited across different *ɛ* clustering thresholds. (F) Effect of success (catch) on the probability of the next spot choice being a revisit across different *ɛ* clustering thresholds. (G) Effect of fish presence when leaving a spot on the probability of the next spot choice being a revisit across different *ɛ* clustering thresholds.

**Extended Data Figure 7.**
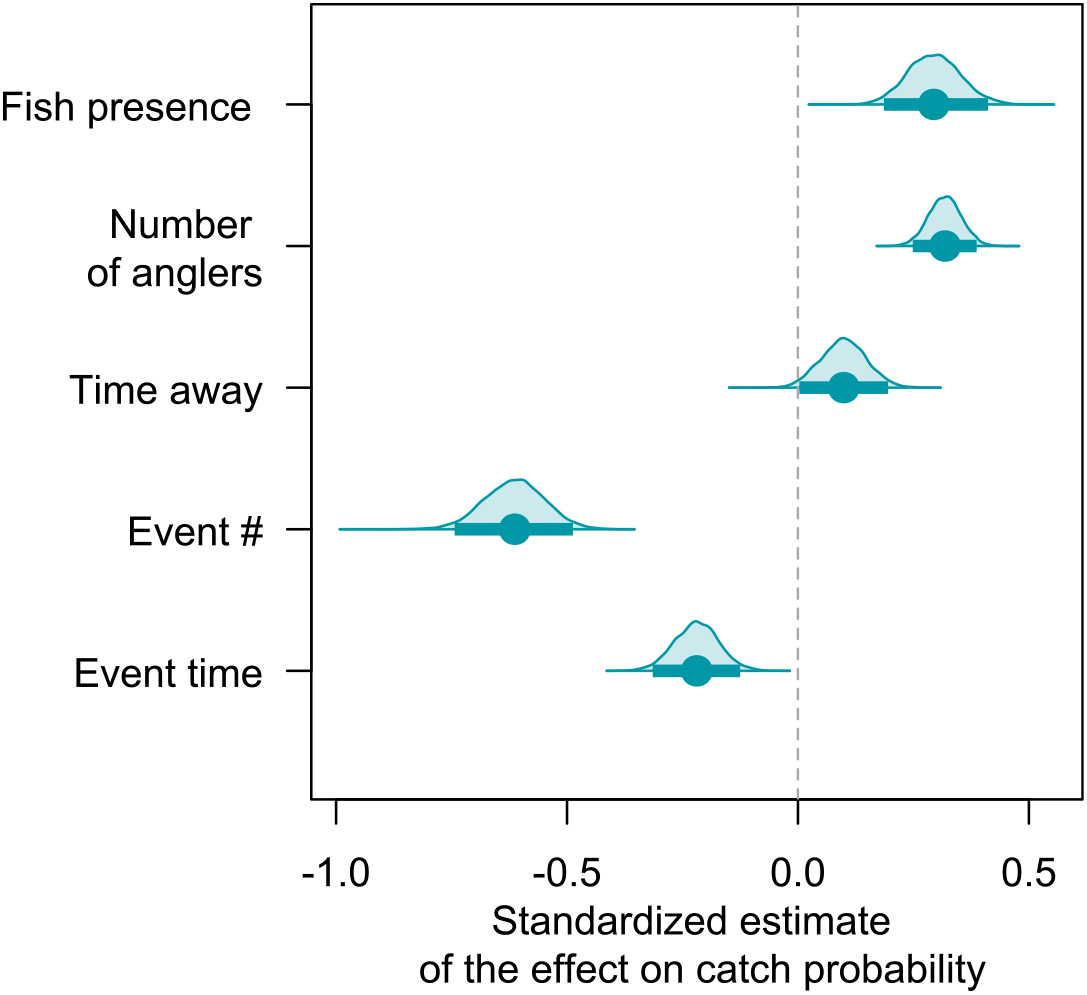
Catch recovery model. Posterior estimates of the model investigating catch recovery when anglers temporarily abandon a spot.

**Extended Data Figure 8.**
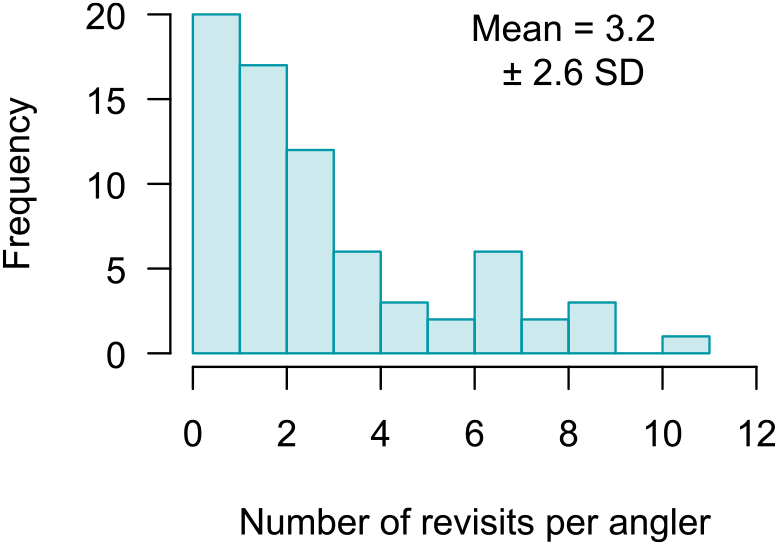
Revisitation rates. Distribution of the number of revisits per angler per competition using an *ɛ* of 10.

## Supplementary Information

**Table S 1.** Sample size and summary statistics per competition.

| Competition | 1 | 2 | 3 | 4 | 5 | 6 | 7 | 8 | 9 | 10 | 11 | 12 | Total |
| --- | --- | --- | --- | --- | --- | --- | --- | --- | --- | --- | --- | --- | --- |
| Date (2024) | 15/03 | 15/03 | 18/03 | 18/03 | 19/03 | 19/03 | 20/03 | 20/03 | 21/03 | 21/03 | 22/03 | 22/03 |  |
| Total angling spots (N) | 155 | 152 | 128 | 173 | 143 | 117 | 140 | 182 | 152 | 165 | 202 | 182 | 1891 |
| Total successful spots (N) | 16 | 17 | 23 | 26 | 42 | 15 | 35 | 39 | 48 | 42 | 51 | 79 | 433 |
| Total fish caught (N) | 56 | 68 | 76 | 191 | 288 | 92 | 262 | 209 | 156 | 142 | 123 | 377 | 2040 |
| Total fish seen on sonar (N) | 2611 | 1628 | 2237 | 2013 | 504 | 887 | 1232 | 3224 | 2019 | 1812 | 1934 | 1559 | 21660 |
| Mean angling spots per trip (N) | 25.8 | 25.3 | 21.3 | 28.8 | 23.8 | 19.5 | 23.3 | 30.3 | 25.3 | 27.5 | 33.7 | 30.3 | 26.2 |
| Mean fish caught per trip (N) | 9.3 | 11.3 | 12.7 | 31.8 | 48 | 15.3 | 43.7 | 34.8 | 26 | 23.7 | 20.5 | 62.8 | 28.3 |
| Mean fish caught per spot (N) | 0.4 | 0.4 | 0.6 | 1.1 | 2 | 0.8 | 1.9 | 1.1 | 1 | 0.9 | 0.6 | 2.1 | 1.1 |
| Mean successful spots per participant (N) | 2.7 | 2.8 | 3.8 | 4.3 | 7 | 2.5 | 5.8 | 6.5 | 8 | 7 | 8.5 | 13.2 | 6 |
| Proportion successful spots per participant (%) | 11.3 | 13.4 | 19.6 | 16.5 | 32.3 | 15.8 | 27.6 | 22.8 | 31.8 | 26.8 | 26.2 | 46.6 | 24.2 |
| Mean angling time at unsuccessful spots (s) | 145.1 | 151.6 | 176.5 | 106.6 | 102.9 | 162.6 | 103.5 | 102.4 | 124.4 | 124.7 | 110.5 | 82 | 124.4 |
| Mean angling time at successful spots (s) | 789.2 | 776.5 | 763.4 | 689.4 | 563 | 1232.2 | 675.7 | 498.7 | 460.4 | 447.9 | 326.1 | 322.8 | 628.8 |
| Mean relocation distance after unsuccessful spots (m) | 24.4 | 18.6 | 19.7 | 16.5 | 21 | 24 | 19.5 | 16.2 | 15.2 | 14.1 | 14.8 | 15.2 | 18.3 |
| Mean relocation distance after successful spots (m) | 11.3 | 7.3 | 7.2 | 9.4 | 15.3 | 10.9 | 13.2 | 8.9 | 6.9 | 11.1 | 9.9 | 11.7 | 10.3 |

**Table S 2.**
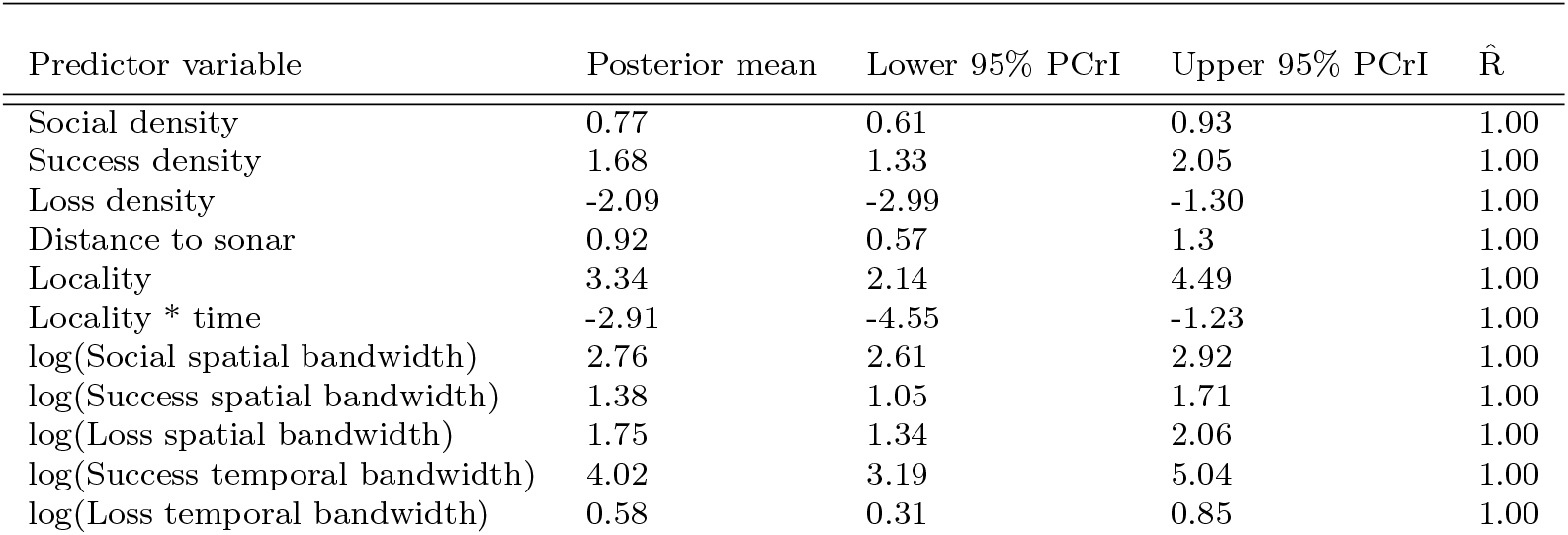
Unstandardized posterior estimates for the spot selection model. PCrI = Posterior credible interval.

**Table S 3.** Posterior estimates for the ecological validation of spot selection model. All explanatory variables are z-scaled. PCrI = Posterior credible interval.

| Predictor variable | Posterior mean | Lower 95% PCrI | Upper 95% PCrI | $\hat{R}$ |
| --- | --- | --- | --- | --- |
| Intercept | 0.44 | 0.07 | 0.79 | 1.00 |
| Social density | 0.29 | 0.24 | 0.35 | 1.00 |
| Success density | 0.09 | 0.04 | 0.14 | 1.00 |
| Loss density | 0.01 | -0.05 | 0.06 | 1.00 |
| Distance to sonar | -0.40 | -0.45 | -0.34 | 1.00 |
| Time | -0.18 | -0.31 | -0.06 | 1.00 |

**Table S 4.** Unstandardized posterior estimates for the spot leaving model. PCrI = Posterior credible interval.

| Predictor variable | Posterior mean | Lower 95% PCrI | Upper 95% PCrI | $\hat{R}$ |
| --- | --- | --- | --- | --- |
| Social density | -0.18 | -0.29 | -0.09 | 1.00 |
| Success density | 0.13 | -0.04 | 0.33 | 1.00 |
| Loss density | 0.43 | -0.06 | 0.89 | 1.00 |
| Time since event | 0.11 | 0.07 | 0.14 | 1.00 |
| Fish catch > 0 | -1.49 | -1.81 | -1.15 | 1.00 |
| Fish presence | -0.33 | -0.55 | -0.12 | 1.00 |
| Distance to sonar | -0.77 | -1.12 | -0.43 | 1.00 |

**Table S 5.** Posterior estimates for the ecological validation of the spot leaving model. Social density, success density, loss density and sonar distance are z-scaled. PCrI = Posterior credible interval.

| Predictor variable | Posterior mean | Lower 95% PCrI | Upper 95% PCrI | $\hat{R}$ |
| --- | --- | --- | --- | --- |
| Social density | 0.32 | 0.30 | 0.34 | 1.00 |
| Success density | 0.03 | 0.00 | 0.05 | 1.00 |
| Loss density | -0.02 | -0.04 | 0.01 | 1.00 |
| Time since event | 0.00 | -0.01 | 0.01 | 1.00 |
| Fish catch > 0 | 0.38 | 0.32 | 0.44 | 1.00 |
| Distance to sonar | -0.31 | -0.34 | -0.29 | 1.00 |

**Table S 6.** Posterior estimates for the angler effect model at unsuccessful spots.

| Predictor variable | Posterior mean | Lower 95% PCrI | Upper 95% PCrI | $\hat{R}$ |
| --- | --- | --- | --- | --- |
| Intercept | -0.004 | -0.006 | -0.002 | 1.00 |
| Time (min) | 0.00006 | -0.0004 | 0.0005 | 1.00 |
| Fish presence at arrival | 0.005 | 0.003 | 0.007 | 1.00 |
| Spot type = visited | 0.005 | 0.003 | 0.008 | 1.00 |
| Distance to sonar | 0.0001 | 0.00007 | 0.0001 | 1.00 |
| Time (min)*Fish presence at arrival | -0.002 | -0.003 | -0.001 | 1.00 |
| Time (min)*Spot type = visited | -0.003 | -0.004 | -0.003 | 1.00 |
| Fish presence at arrival*Spot type = visited | -0.005 | -0.008 | -0.001 | 1.00 |
| Time (min)*Fish presence at arrival*Spot type = visited | 0.002 | 0.0009 | 0.003 | 1.00 |

**Table S 7.** Posterior estimates for the angler effect model at successful spots. PCrI = Posterior credible interval.

| Predictor variable | Posterior mean | Lower 95% PCrI | Upper 95% PCrI | $\hat{R}$ |
| --- | --- | --- | --- | --- |
| Intercept | 0.03 | 0.01 | 0.04 | 1.00 |
| Time (min) | 0.003 | 0.002 | 0.004 | 1.00 |
| Fish presence at arrival | 0.0001 | -0.01 | 0.01 | 1.00 |
| Spot type = visited | -0.03 | -0.04 | -0.01 | 1.00 |
| Distance to sonar | -0.001 | -0.001 | -0.0009 | 1.00 |
| Time (min)*Fish presence at arrival | -0.006 | -0.007 | -0.005 | 1.00 |
| Time (min)*Spot type = visited | 0.003 | 0.002 | 0.004 | 1.00 |
| Fish presence at arrival*Spot type = visited | 0.03 | 0.02 | 0.05 | 1.00 |
| Time (min)*Fish presence at arrival*Spot type = visited | -0.003 | -0.004 | -0.001 | 1.00 |

**Table S 8.**
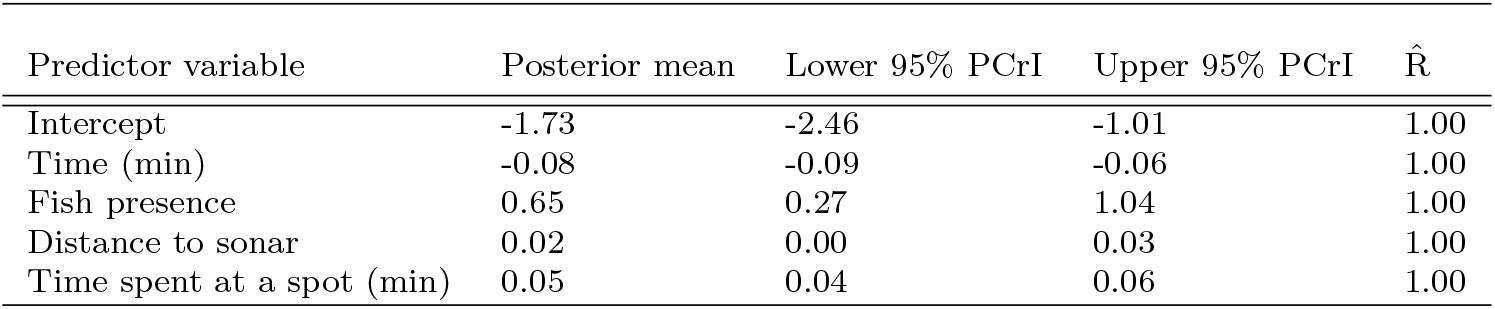
Posterior estimates for the model investigating how catch likelihood varies over time and fish presence. PCrI = Posterior credible interval.

**Table S 9.** Posterior estimates for the model investigating catch recovery in the absence of anglers. PCrI = Posterior credible interval.

| Predictor variable | Posterior mean | Lower 95% PCrI | Upper 95% PCrI | $\hat{R}$ |
| --- | --- | --- | --- | --- |
| Intercept | -1.85 | -2.26 | -1.45 | 1.00 |
| Time in event | -0.01 | -0.02 | -0.007 | 1.00 |
| Event number | -0.29 | -0.35 | -0.23 | 1.00 |
| Time away | 0.001 | 0.00003 | 0.002 | 1.00 |
| Number of anglers | 0.67 | 0.52 | 0.81 | 1.00 |
| Fish presence | 0.98 | 0.62 | 1.38 | 1.00 |

**Table S 10.** Posterior estimates for the model investigating whether success at a spot predicts the likelihood of returning to that same spot later. PCrI = Posterior credible interval.

| Predictor variable | Posterior mean | Lower 95% PCrI | Upper 95% PCrI | $\hat{R}$ |
| --- | --- | --- | --- | --- |
| Intercept | 0.88 | 0.45 | 1.33 | 1.00 |
| Success | 0.83 | 0.58 | 1.08 | 1.00 |
| Time at which the spot is left | -0.03 | -0.03 | -0.02 | 1.00 |

**Table S 11.**
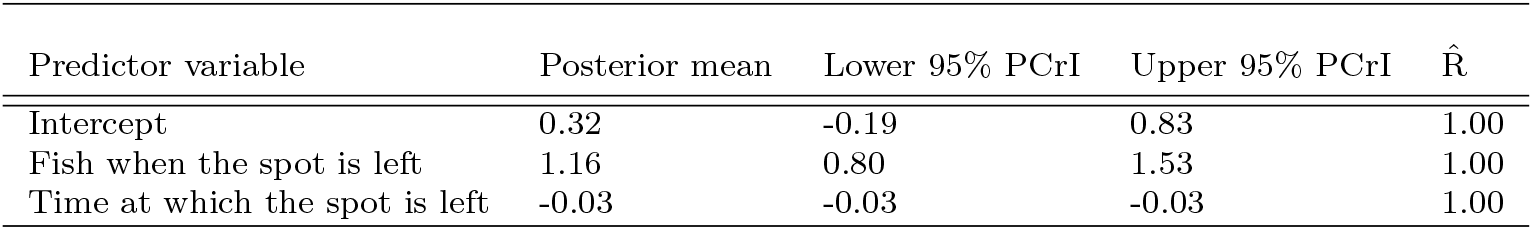
Posterior estimates for the model investigating whether fish presence when a spot is left predicts the likelihood of returning to that same spot later. PCrI = Posterior credible interval.

**Table S 12.** Posterior estimates for the model investigating whether a spot choice is a revisit given the success at the previous spot. PCrI = Posterior credible interval.

| Predictor variable | Posterior mean | Lower 95% PCrI | Upper 95% PCrI | $\hat{R}$ |
| --- | --- | --- | --- | --- |
| Intercept | -2.40 | -2.86 | -1.96 | 1.00 |
| Previous success | -1.44 | -2.01 | -0.93 | 1.00 |
| Time at which the spot choice is made | 0.00 | -0.00 | 0.01 | 1.00 |
| Spot number | 0.03 | 0.00 | 0.05 | 1.00 |

**Table S 13.** Posterior estimates for the model investigating whether a spot choice is a revisit given the fish presence at the previous spot. PCrI = Posterior credible interval.

| Predictor variable | Posterior mean | Lower 95% PCrI | Upper 95% PCrI | $\hat{R}$ |
| --- | --- | --- | --- | --- |
| Intercept | -2.49 | -3.03 | -1.97 | 1.00 |
| Previous fish presence | -0.24 | -0.68 | 0.21 | 1.00 |
| Time at which the spot choice is made | 0.00 | -0.00 | 0.01 | 1.00 |
| Spot number | 0.03 | 0.00 | 0.05 | 1.00 |

**Table S 14.** Optimized hyperparameters of the GP models for each competition.

| Competition | Constant | Space | Time | $\alpha$ | MSE |
| --- | --- | --- | --- | --- | --- |
| Sompalampi March 15 am | 1.7 | 5.3 | 16.97 | $1.16 \times 10^{-5}$ | $9.69 \times 10^{-15}$ |
| Sompalampi March 15 pm | 1.2 | 7.1 | 15.67 | $1.16 \times 10^{-5}$ | $8.23 \times 10^{-15}$ |
| Sompalampi March 18 am | 1.7 | 5.3 | 16.97 | $1.16 \times 10^{-5}$ | $6.37 \times 10^{-15}$ |
| Sompalampi March 18 pm | 1.7 | 5.3 | 16.97 | $1.16 \times 10^{-5}$ | $5.53 \times 10^{-15}$ |
| Kuorinka March 19 am | 1.2 | 7.1 | 15.67 | $1.16 \times 10^{-5}$ | $1.67 \times 10^{-14}$ |
| Kuorinka March 19 pm | 1.7 | 5.3 | 16.99 | $1.16 \times 10^{-5}$ | $1.33 \times 10^{-14}$ |
| Kuorinka March 20 am | 1.7 | 5.3 | 16.97 | $1.16 \times 10^{-5}$ | $9.33 \times 10^{-15}$ |
| Kuorinka March 20 pm | 1.2 | 7.1 | 15.67 | $1.16 \times 10^{-5}$ | $8.54 \times 10^{-15}$ |
| Pitkanen March 21 am | 1.7 | 5.3 | 16.97 | $1.16 \times 10^{-5}$ | $5.69 \times 10^{-15}$ |
| Pitkanen March 21 pm | 1.2 | 7.1 | 15.67 | $1.16 \times 10^{-5}$ | $4.91 \times 10^{-15}$ |
| Pitkanen March 22 am | 1.2 | 7.1 | 15.67 | $1.16 \times 10^{-5}$ | $4.05 \times 10^{-15}$ |
| Pitkanen March 22 pm | 1.7 | 5.3 | 16.97 | $1.16 \times 10^{-5}$ | $1.07 \times 10^{-14}$ |

